# Comprehensive Proteomic Characterization of the Intra-Golgi Trafficking Intermediates

**DOI:** 10.1101/2024.10.25.620336

**Authors:** Farhana Taher Sumya, Walter S. Aragon-Ramirez, Vladimir V Lupashin

**Affiliations:** University of Arkansas for Medical Sciences, Department of Physiology and Cell Biology, Little Rock, Arkansas, US

**Keywords:** Golgi, vesicular trafficking, glycosylation, COG complex, vesicle tethering, SNARE, vesicular coat, degron, mass-spectrometry

## Abstract

Intracellular trafficking relies on small vesicular intermediates, though their specific role in Golgi function is still debated. To clarify this, we induced acute dysfunction of the Conserved Oligomeric Golgi (COG) complex and analyzed vesicles from cis, medial, and trans-Golgi compartments. Proteomic analysis of Golgi-derived vesicles from wild-type cells revealed distinct molecular profiles, indicating a robust recycling system for Golgi proteins. Notably, these vesicles retained various vesicular coats, while COG depletion accelerated uncoating. The increased overlap in molecular profiles with COG depletion suggests that persistent defects in vesicle tethering disrupt intra-Golgi sorting.

Our findings reveal that the entire Golgi glycosylation machinery recycles within vesicles in a COG-dependent manner, whereas secretory and ER-Golgi trafficking proteins were not enriched. These results support a model in which the COG complex orchestrates multi-step recycling of glycosylation machinery, coordinated by specific Golgi coats, tethers, Rabs, and SNAREs.

## Introduction

Membrane trafficking is essential in all eukaryotic cells. Over 30% of cellular proteins are modified within the secretory pathway, which depends on the correct localization of modifying enzymes and the effective function of the membrane trafficking machinery(D’Souza et al., 2020; Wallin & von Heijne, 1998). Proteins in this pathway can be either soluble or transmembrane and may be directed to various destinations, internally or externally, depending on the specific protein. Vesicular membrane trafficking within cells relies on a delicate balance of anterograde and retrograde transport to ensure delivery of proteins and lipids to their correct destinations and to maintain organelle homeostasis. The Golgi apparatus serves as the cell’s primary packaging, sorting, and processing center (Füllekrug & Nilsson, 1998; Kienzle & von Blume, 2014; Maccioni et al., 2002; Munro, 1998; Polishchuk & Mironov, 2004). Glycosylation, a major modification of cargo is primarily carried out by the Golgi glycosylation machinery (Stanley, 2011). Several models have been proposed to explain how Golgi resident proteins and enzymes are sorted and retained (Banfield, 2011; Emr et al., 2009; Glick & Nakano, 2009; Nilsson et al., 2009; Pantazopoulou & Glick, 2019; Rothman & Wieland, 1996; Sahu et al., 2022; Tu & Banfield, 2010; Welch & Munro, 2019, 2019). All these models include vesicle-mediated trafficking between Golgi cisternae, but the details of intra-Golgi vesicle trafficking are poorly understood. The vesicular trafficking machinery consists of several distinct modules responsible for driving vesicle budding from a donor compartment, its subsequent transport, tethering, and fusion with the acceptor compartment (Bonifacino & Glick, 2004; Cai et al., 2007; Cottam & Ungar, 2012; Park et al., 2021). Vesicle tethering is achieved by both coiled-coil and multi-subunit tethering complex (MTC) tethers, followed by vesicle fusion with a specific Golgi subcompartment in a SNARE-dependent reaction (Arab et al., 2024; Blackburn et al., 2019; Bonifacino & Glick, 2004; Gillingham & Munro, 2016; Stanton & Hughson, 2023; Willett, Ungar, et al., 2013). The Conserved Oligomeric Golgi (COG) complex plays a crucial role in intra-Golgi retrograde trafficking by interacting with various cellular components of the vesicle docking/fusion machinery through specific interaction sites on its subunits (D’Souza et al., 2020; Laufman et al., 2013; Miller et al., 2013; Stanton & Hughson, 2023; Ungar et al., 2002, 2006; Willett, Kudlyk, et al., 2013). In humans, malfunction of COG can lead to global glycosylation defects known as COG-related congenital disorders of glycosylation (COG-CDGs) (D’Souza et al., 2020; Foulquier, 2009; Sumya et al., 2021). Previous studies conducted by our lab and others have shown that malfunction or RNAi-induced depletion of COG leads to the accumulation of non-tethered COG complex-dependent (CCD) vesicles (Cottam et al., 2014; Wuestehube et al., 1996; Zolov & Lupashin, 2005). The accumulation of CCD vesicles is transient (Shestakova et al., 2006), which created technical challenges for their isolation and characterization. Moreover, the extent of intra-Golgi recycling of Golgi resident proteins, specifically the Golgi glycosylation machinery, is still a matter of fierce debate. It is not clear how intimately COG deficiency is connected to the degradation of Golgi enzymes and how this degradation is related to vesicular trafficking. The introduction of a degron-driven COG4 degradation system has enabled the acute and consistent accumulation of CCD vesicles (Sumya, Pokrovskaya, D’Souza, et al., 2023; Sumya, Pokrovskaya, & Lupashin, 2023). In this study, we combined degron-assisted COG4 degradation with vesicle immunoprecipitation and data-independent acquisition (DIA) mass spectrometry to comprehensively characterize intra-Golgi trafficking intermediates. This approach allowed us to evaluate the overall dependence of intra-Golgi trafficking on COG function and provided deeper insight into the composition and function of COG-dependent trafficking intermediates.

## Materials and Methods

### Cell Culture and Auxin Treatment

hTERT RPE1 (Retinal Pigment Epithelial) cells were purchased from ATCC. hTERT RPE1 COG4 KO expressing OsTIR1-9myc (RPE1-COG4KO-OsTIR1) or co-expressing OsTIR1 and COG4-mAID-mCherry (RPE1-COG4-mAID-mCherry) cells were described previously (Sumya, Pokrovskaya, D’Souza, et al., 2023). Cells were cultured in Dulbecco’s Modified Eagle’s Medium (DMEM) containing Nutrient mixture F-12 (DMEM/F12, Corning #10–092-CV) supplemented with 10% Fetal Bovine Serum (Atlas Biologicals, CF-0500-A). Cells were incubated in a 37°C incubator with 5% CO_2_ and 90% humidity.

For rapid COG4 degradation, a stock solution of 0.5 M indole-3-acetic acid sodium salt (auxin, IAA, Sigma # I5148) was prepared in water and stored in a frozen aliquot. Time course treatment of cells was performed with 500 µM IAA for 2 hours at 37°C. The RPE1 WT cells with auxin treatment (first experiment) and the hTERT RPE1 COG4 KO co-expressing OsTIR1 and COG4-mAID cells without auxin treatment (second experiment) were considered as controls.

### Construction of COG4-mAID-3myc plasmids

To produce COG4-mAID-3myc in pENTRA1A, mAID portion was first amplified by PCR from COG4-mAID-mCherry in pENTRA1A (Sumya, Pokrovskaya, D’Souza, et al., 2023) using primers 5′-GCCTGGGTACCGGATCCGGTGCAG-3′, and 5′-GGCGGGTACCTTTATACATCCTCAAATCGAT-3′ following KpnI digestion and ligation of PCR fragment with similarly digested COG4-3myc in pEntra1A (Sumya et al., 2021).

The COG4-mAID-3myc in pEntra1A was recombined into the pLenti COG4_pr_ Neo DEST (as referenced in (Khakurel et al., 2021; Sumya et al., 2021; Sumya, Pokrovskaya, D’Souza, et al., 2023) using the Gateway LR Clonase II Enzyme Mix (Thermo Fisher). The resulting COG4-mAID-3myc pLenti plasmid was then transformed into Stbl3 competent cells following the manufacturer’s instructions. DNA extraction was performed using the QIAprep Spin Miniprep DNA Extraction Kit. COG4-mAID-3myc pLenti clones were verified through restriction analysis. The expression of COG4-mAID-3myc was confirmed by transfecting HEK293T cells with selected COG4-mAID-3myc pLenti plasmids, followed by western blot analysis of total cell lysates using COG4 and myc antibodies.

### Production of COG4-mAID-myc lentivirus and RPE1-COG4-mAID-3myc stable cell line

To produce lentiviral particles, equal amounts of the lentiviral packaging plasmids pMD2.G (gift from Didier Trono; Addgene plasmid #12259, http://n2t.net/addgene: 12259, RRID: Addgene_12259), pRSV-Rev, pMDLg/pRRE (Dull et al., 1998), and COG4-mAID-3myc pLenti were mixed and transfected into HEK293FT cells using Lipofectamine 3000 following the manufacturer’s protocol. The transfected cells were placed in serum-reduced Opti-MEM supplemented with 25 μM Chloroquine and 1× GlutaMAX. The next day, the medium was changed to Opti-MEM supplemented with 1× GlutaMAX. The medium was collected after 72 hours of transfection, t, and cell debris was removed by centrifugation at 600×Lg for 10Lmin. The supernatant was filtered through a 0.45 μM polyethersulfone (PES) membrane filter, and the lentiviral medium was stored at 4°C overnight or divided into aliquots, snap-frozen in liquid nitrogen, and stored at −80°C.

RPE1-COG4KO-OsTIR1 cells (Sumya, Pokrovskaya, D’Souza, et al., 2023) were seeded in two wells of a 6-well plate with complete media to achieve 90% confluency the following day. One well was designated as a control for antibiotic selection. The next day, the cells were transduced with 500 μl of lentiviral supernatant. After 48 hours post-transduction, the lentiviral media was replaced with cell growth media containing G418 (500 μg/mL final concentration, selection dose). Following 48 hours of selection, the media was switched to complete media containing 200Lμg/mL of G418 (maintenance dose). The cells were then cultured at 37°C and 5% CO_2_ for 48Lhours. Following G418 selection, single-cell clones were isolated into 96-well plates by serial dilution. The cells were allowed to grow for two weeks, collected by trypsin treatment, and each colony was expanded into a 12 well plate (4 cm^2^) with complete media containing G418. Subsequently, WB and IF analyses were conducted to identify clones with COG4-mAID-3myc expression. Clones demonstrating uniform expression of COG4-mAID-3myc (RPE1-COG4-mAID-3myc) were transferred to 10-cm dishes, and aliquots were cryopreserved in 2× freezing medium (80% FBS with 20% DMSO) mixed with growth medium.

### Immunoprecipitation of Golgi-derived vesicles

Cells were cultured in 15 cm dishes, until they reached 90% confluency, rinsed with PBS and harvested by trypsinization, followed by centrifugation at 400×Lg for 5 min. The cell pellet was then resuspended in 1.5 ml of cell collection solution (0.25 M sucrose in PBS) and centrifuged at 400×Lg for 5 min. Subsequently, the pellet was resuspended in 1.5 ml of a hypotonic lysis solution (20LmM HEPES pHL7.2, containing protein inhibitor cocktail and 1 mM PMSF) and passed through a 25LG needle 20 times to disrupt the cells. The efficiency of cell lysis was assessed using phase-contrast microscopy. Following this, KCL (to a final concentration of 150LmM) and EDTA (to a final concentration of 2 mM) were added. Unlysed cells and cell nuclei were separated by centrifugation at 1000×Lg. The postnuclear supernatant (PNS) was then transferred to a 1.5 ml Beckman tube (#357488), and the Golgi-enriched fraction was sedimented at 30L000×Lg for 10 min. The supernatant (S30) was transferred into a new Beckman tube. Samples from each fraction were prepared for WB analysis, while the remaining samples were utilized for IP or MS analysis.

In the initial step of the first vesicle IP experiment, the S30 supernatant was combined with an affinity-purified anti-Giantin antibody (1.33 µg/mL) and left to incubate at room temperature on a rotating platform for one hour. Following this, 30 µl of Dyna Protein G magnetic beads (ThermoFisher Scientific #10004D) were added, and the mixture was rotated at room temperature for an additional hour. Subsequently, the magnetic beads with attached Giantin-positive membrane were isolated using the DynaMag^TM^-2 (Magnetic particle concentrator, ThermoFisher), washed three times in a wash buffer consisting of 20LmM HEPES pHL7.2, 1LmM PMSF, 150 mM KCL, 2 mM EDTA. Proteins bound to the beads (G-vIP) were extracted by adding 2X sample buffer with 10% β-mercaptoethanol and heated at 95°C in a heat block for 5 minutes. Next, the unbound material, flow through (FT) from the first vesicle IP (S30-G-FT) was successively exposed to affinity-purified anti-golgin-84 antibody for one hour to produce g84–vIP and S30-g84-FT. Finally, S30-g84-FT was incubated with affinity-purified anti-GS15 antibody for two hours, producing GS15-vIP and ST-FT. In the second vesicle IP, the vesicle isolation procedure was identical to the first experiment except the sequence of the antibody precipitations. In this experiment the S30 was sequentially incubated with STX5, Giantin, VAMP7, and GS15 antibodies for one hour, except for GS15 (two hours).

### Western Blot Analysis

Protein samples (10-20 µg) were loaded onto either a Bio-Rad (4-15%) or Genescript (8-16%) gradient gel. The proteins were then transferred to a 0.2 µm nitrocellulose blotting membrane (Amersham^TM^ Protran^TM^) using the Thermo Scientific Pierce G2 Fast Blotter. Afterward, the membranes were washed in PBS, blocked in Bio-Rad blocking buffer for 20 minutes, and incubated with primary antibodies for 1 hour at room temperature or overnight at 4°C. Following this, the membranes were washed with PBS and incubated with secondary fluorescently-tagged antibodies diluted in Bio-Rad blocking buffer for 1 hour. The primary and secondary antibodies used in the study are listed in Table 1. The blots were then washed with PBS and imaged using the Odyssey Imaging System. Images were processed using the LI-COR Image Studio software.

**Table 1.**
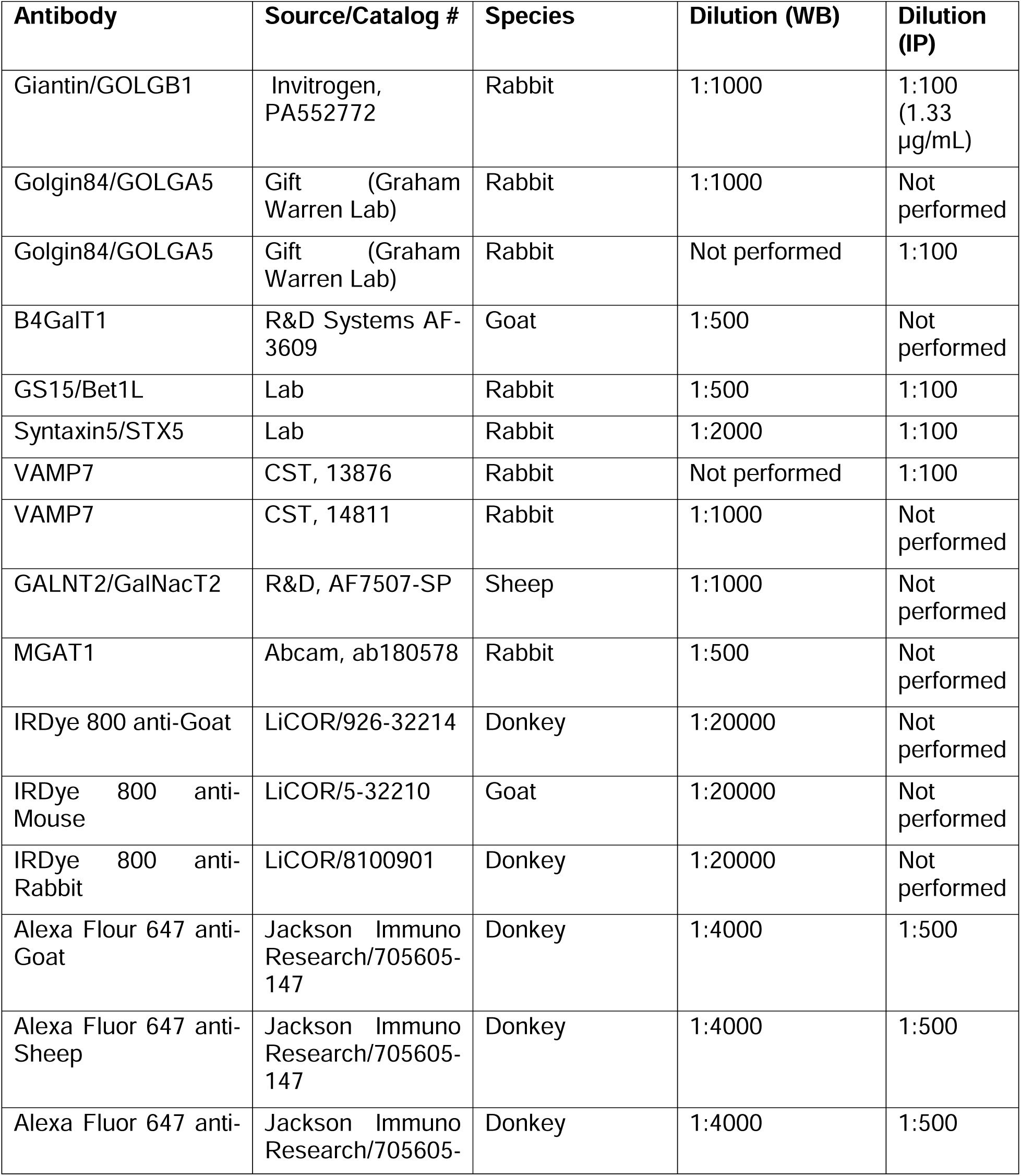
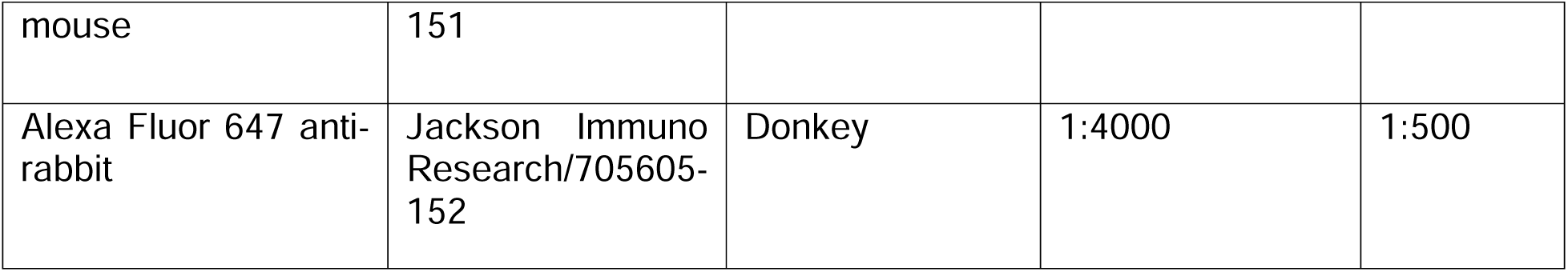
List of antibodies.

### Proteomic analysis using Orbitrap Exploris DIA

The vesicle IP was performed as described above. For Mass Spectrometry (MS), the Protein-G magnetic beads were washed three more times in a wash buffer to remove excess detergent before eluting.

Purified proteins were reduced, alkylated, and digested on-bead using a filter-aided sample preparation (Wiśniewski et al., 2009) with sequencing grade-modified porcine trypsin (Promega). Tryptic peptides were then separated by reverse phase XSelect CSH C18 2.5 um resin (Waters) on an in-line 150 x 0.075 mm column using the UltiMate 3000 RSLCnano system (Thermo). Peptides were eluted using a 60 min gradient from 98:2 to 65:35 buffer A:B ratio.

Buffer A = 0.1% formic acid, 0.5% acetonitrile

Buffer B = 0.1% formic acid, 99.9% acetonitrile

Eluted peptides were ionized by electrospray (2.2 kV) followed by mass spectrometric analysis on an Orbitrap Exploris 480 mass spectrometer (Thermo). To assemble a chromatogram library, six gas-phase fractions were acquired on the Orbitrap Exploris with 4 m/z DIA spectra (4 m/z precursor isolation windows at 30,000 resolution, normalized AGC target 100%, maximum inject time 66 ms) using a staggered window pattern from narrow mass ranges with optimized window placements. Precursor spectra were acquired after each DIA duty cycle, spanning the m/z range of the gas-phase fraction (i.e. 496-602 m/z, 60,000 resolution, normalized AGC target 100%, maximum injection time 50 ms). For wide-window acquisitions, the Orbitrap Exploris was configured to acquire a precursor scan (385-1015 m/z, 60,000 resolution, normalized AGC target 100%, maximum injection time 50 ms) followed by 50x 12 m/z DIA spectra (12 m/z precursor isolation windows at 15,000 resolution, normalized AGC target 100%, maximum injection time 33 ms) using a staggered window pattern with optimized window placements. Precursor spectra were acquired after each DIA duty cycle.

Following data acquisition, data were searched using an empirically corrected library against the UniProt *Homo sapiens* database (April 2022), and a quantitative analysis was performed to obtain a comprehensive proteomic profile. Proteins were identified and quantified using EncyclopeDIA (Searle et al., 2018) and visualized with Scaffold DIA using 1% false discovery thresholds at both the protein and peptide level. Protein MS2 exclusive intensity values were assessed for quality using ProteiNorm (Graw et al., 2020). The data was normalized using cyclic loess (Ritchie et al., 2015) and analyzed using proteoDA to perform statistical analysis using Linear Models for Microarray Data (limma) with empirical Bayes (eBayes) smoothing to the standard errors (Ritchie et al., 2015; Thurman et al., 2023). Proteins with an FDR adjusted p-value < 0.05 and a fold change >2 were considered significant.

### Proteomic analysis using Orbitrap Exploris 480 DIA

The vesicle IP was performed as described above. For MS, the Protein-G magnetic beads were washed three more times in wash buffer before eluting to remove excess detergent.

Protein samples were reduced, alkylated, and digested on-bead using a filter-aided sample preparation (Wiśniewski et al., 2009) with sequencing grade-modified porcine trypsin (Promega). Tryptic peptides were trapped and eluted on 3.5um CSH C18 resin (Waters) (4mm x 75um) then separated by reverse phase XSelect CSH C18 2.5 um resin (Waters) on an in-line 150 x 0.075 mm column using the UltiMate 3000 RSLCnano system (Thermo). Peptides were eluted at a flow rate of 0.300uL/min using a 60 min gradient from 98% Buffer A:2% Buffer B to 95:5 at 2.0 minutes to 80:20 at 39.0 minutes to 60:40 at 48.0 minutes to 10:90 at 49.0 minutes and hold until 53.0 minutes and then equilibrated back to 98:2 at 53.1 minutes until 60 minutes.

Buffer A = 0.1% formic acid, 0.5% acetonitrile

Buffer B = 0.1% formic acid, 99.9% acetonitrile

Eluted peptides were ionized by electrospray (2.4 kV) through a heated capillary (275°C) followed by data collection on an Orbitrap Exploris 480 mass spectrometer (Thermo Scientific). Precursor spectra were acquired with a scan from 385-1015 Th at a resolution set to 60,000 with 100% AGC, max time of 50 msec, and an RF parameter at 40%. DIA was configured on the Orbitrap 480 to acquire 50 x 12 Th isolation windows at 15,000 resolution, normalized AGC target 500%, maximum injection time 40 ms). A second DIA was acquired in a staggered window (12 Th) pattern with optimized window placements.

Following data acquisition, data were searched using Spectronaut (Biognosys version 18.6) against the UniProt *Homo sapiens* database (January, 2024) using the directDIA method with an identification precursor and protein q-value cutoff of 1%, generate decoys set to true, the protein inference workflow set to maxLFQ, inference algorithm set to IDPicker, quantity level set to MS2, cross-run normalization set to false, and the protein grouping quantification set to median peptide and precursor quantity. Protein MS2 intensity values were assessed for quality using ProteiNorm (Graw et al., 2020). The data was normalized using RLR (robust linear regression) and analyzed using proteoDA to perform statistical analysis using Linear Models for Microarray Data (limma) with empirical Bayes (eBayes) smoothing to the standard errors (Ritchie et al., 2015; Thurman et al., 2023). Proteins with an FDR (False Discovery Rate) adjusted p-value <0.05 and a fold change >2 were considered significant.

### CCD vesicle re-routing to Golgin-84-decorated mitochondria

The Golgin84-HA-OMP DNA construct contains human GOLGA5 (residues 1-698) fused with HA epitope and C-terminal transmembrane domain of OMP25 (aa 110-145) was synthesized by GenScript and subcloned into pcDNA3.1. A similar construct without the GOLGA5 sequence was used as a control.

COG4-mAID cells were cultivated on 12-mm round coverslips until they reached 80%-90% confluency. Subsequently, they were transiently transfected with Golgin-84-HA-OMP using Lipofectamine 3000 as per the manufacturer’s protocol. After 24 hours of transfection, the cells were treated with IAA for two hours to accumulate CCD vesicles. Control and IAA-treated cells were fixed with freshly prepared 4% paraformaldehyde (PFA) diluted in PBS for 15 minutes at room temperature. The cells were permeabilized with 0.1% Triton X-100 for 1 minute, followed by treatment with 50 mM ammonium chloride for 5 minutes, and then washed twice with PBS. Blocking was performed by incubating the cells twice for 10 minutes each in 1% BSA and 0.1% saponin in PBS. Subsequently, the cells were incubated with the primary antibody (diluted in 1% cold fish gelatin and 0.1% saponin in PBS) for 45 minutes, washed, and then incubated with fluorescently conjugated secondary antibodies diluted in the antibody buffer for 30 minutes. After washing the cells four times with PBS, the coverslips were dipped in PBS and water 10 times each and then mounted on glass microscope slides using Prolong® Gold antifade reagent (Life Technologies). Finally, the cells were imaged using a 63X oil 1.4 numerical aperture (NA) objective of an LSM880 Zeiss Laser inverted microscope with Airyscan using ZEN software.

### Statistical Analysis

All mass spectrometry results are based on at least four biological replicates. The Western blot images represent two repeats and were quantified using LI-COR Image Studio software. Error bars in all graphs indicate standard deviation. The GO enrichment analysis was conducted using the Shiny GO 0.81 website, and Venn diagrams were created using the “Bioinformatics and Evolutionary Genomics” website.

## Results

### Isolation of Golgi-derived vesicles from control and COG complex–deficient cells

Previous studies reported the accumulation of heterogenous non-tethered COG complex-dependent (CCD) vesicles in cells acutely depleted for COG3 (Shestakova et al., 2006; Zolov & Lupashin, 2005) or COG4 (Sumya, Pokrovskaya, D’Souza, et al., 2023) of the COG complex subunits. Since CCD vesicles accumulate several Golgi resident proteins with large cytoplasmic domains, native immunocapturing was utilized for purification and characterization of the trafficking intermediates. To obtain crude vesicle fraction, both control and COG4-depleted hTERT-RPE1 cells were first disrupted by a gentle mechanical procedure (Sumya, Pokrovskaya, D’Souza, et al., 2023) followed by differential centrifugation to remove large membranes (nucleus, mitochondria, ER, plasma membrane, and Golgi). Next, Golgi-derived vesicles were isolated from the S30 fraction (total vesicle fraction) by native membrane immunoprecipitation (IP) using affinity-purified antibodies to Golgi-resident transmembrane proteins and Protein-G magnetic beads. Cis-medial resident Giantin/GOLGB1 (Linstedt & Hauri, 1993; Sönnichsen et al., 1998), medial Golgi resident golgin-84/GOLGA5 (Bascom et al., 1999; Diao et al., 2003; Satoh et al., 2003) and trans-Golgi resident GS15/BET1L (Volchuk et al., 2004) were chosen as “handles” for isolation of different vesicle populations **(Figure 1A)**. These transmembrane (TM) proteins were previously found in Golgi-derived vesicles and significantly redistributed to the CCD vesicle fraction upon COG dysfunction (Sohda et al., 2010; Sönnichsen et al., 1998; Sumya, Pokrovskaya, D’Souza, et al., 2023; Zolov & Lupashin, 2005). The vesicle IPs (vIP from control cells and CCDvIP from COG4-depleted cells) were performed consecutively. The first precipitation was done with giantin antibodies (G-vIP and G-CCDvIP). Flow-through from the Giantin-IP was precipitated with anti-golgin84 (g84-vIP and g84-CCDvIP), and the golgin84-IP flow-through was used to precipitate GS15-positive vesicles (GS15-vIP and GS15-CCDvIP) **(Figure 1A)**. WB analysis revealed the enrichment of Giantin, golgin-84, and GS15 in the Giantin-IP, golgin-84, and GS15-IP, respectively, validating the approach **(Figure 1B)**. Some Giantin was also detected in golgin-84 vesicles isolated from COG4-depleted cells, and some golgin-84 signal was found in GS15 IPs, suggesting that the immunoisolated vesicle populations are partially mixed and may not represent distinct entities entirely. This was expected, as the IP time was intentionally shortened to minimize protein and membrane degradation. However, the immunoisolated vesicle populations remain significantly distinct, and follow-up proteomic analysis (see below) confirmed this distinction.

**Figure 1.**
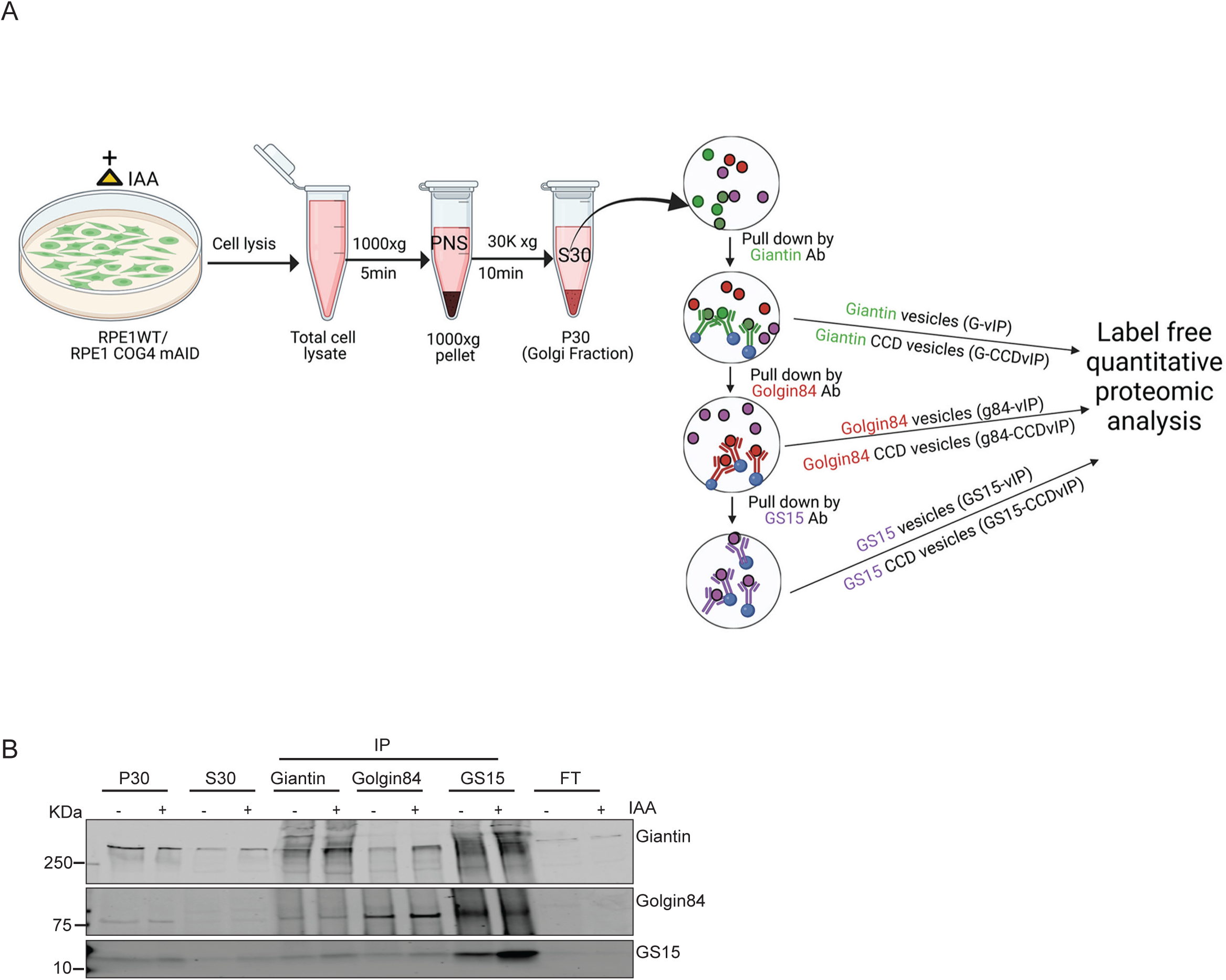
Immunoisolation of Golgi-derived vesicles. **A)** Schematic diagram showing immune-isolation of vesicles from RPE1 WT and RPE1 COG4 KO cell line expressing COG4-mAID-mCherry and OsTIR1-9myc (RPE1 COG4-mAID cells) using Giantin, Golgin-84 and GS15 antibodies after 2 hours of IAA treatment for Mass Spectrometric Analysis. The vesicles isolated from control cells are named G-vIP, g84-vIP, and GS15-vIP, and the vesicles isolated from acute COG-depleted cells are called G-CCDvIP, g84-CCDvIP, and GS15-CCDvIP, respectively. **B)** WB analysis of Giantin, Golgin-84, and GS15 proteins in Golgi fraction (P30), a soluble fraction (S30), Giantin, Golgin-84, and GS15 pull-down vesicles before and after IAA treatment.

To account for potential trafficking abnormalities associated with the usage of tagged COG4, a second vesicle IP experiment was conducted. In this setup, vesicles were immunoprecipitated from the same cell line, either untreated or treated with IAA to induce COG4 degradation **(Supplementary Figure 1A).** To eliminate potential vesicle contamination from small fragments of Golgi cisternae that may result from mechanical disruption, the S30 fraction was pre-cleared with STX5 IP in the second experiment. While STX5 interacts with the COG complex (Shestakova et al., 2007; Suvorova et al., 2002), it does not partition into CCD vesicles (Sumya, Pokrovskaya, D’Souza, et al., 2023). We were out of custom-made golgin-84 antibodies for IP, and commercial antibodies are unavailable; therefore, in the second experiment, commercial affi-pure antibodies to VAMP7 were used. We have shown previously that VAMP7 is a part of STX5/SNAP29/VAMP7 Golgi SNARE complex, and that anti-VAMP7 antibody are suitable for native IP (D’Souza et al., 2023). Additionally, the first round of vesicle IP indicated that VAMP7 is a COG-sensitive protein that is significantly increased in abundance in G-CCDvIP and GS15-CCDvIP **(Table S1)**. The second vesicle IP was evaluated by label-free MS analysis using more sensitive detection tools (see Materials and Methods), which allows the detection of more proteins than the first experiment **(Table S2)**. Similarly to the first experiment, vesicle IPs were validated by WB analysis **(Supplementary Figure 1B)**.

### Exploring the proteome of Golgi-derived vesicles

Label-free quantitative mass spectrometric analysis of immunoprecipitated vesicles resulted in the identification of 2172 proteins **(Table S1)**. Among of these proteins 417 were previously detected in the “Golgi set” (Fasimoye et al., 2023), and an additional 16 are potential Golgi resident proteins. Notably, the degree of recovery of Golgi resident proteins in vesicle-IPs was comparable to the number of resident proteins identified in recently published Golgi-IP (Fasimoye et al., 2023), indicating the high efficiency of vesicle-IPs, validating our approach. Results of the second round of vesicle IP included 5638 proteins (Table S2), with 739 of them found in the “Golgi set” (Fasimoye et al., 2023), of which 37 are putative Golgi proteins. The first vesicle IP resulted in the identification of 62 members of the Golgi glycosylation machinery, 146 other transmembrane (TM), 54 lumenal/secretory, and 171 peripheral Golgi proteins. The second set of vesicle IP detected 110 members of the Golgi glycosylation machinery, 296 other TM, 89 lumenal/secretory, and 281 peripheral Golgi proteins **(Table S5)**. For further analysis, we focused on proteins that were significantly enriched at least 2-fold (logFC > 1) in the vesicle IP compared to the crude vesicle fraction (S30). First, GO enrichment analysis was performed on 696 proteins in G-vIP, 658 proteins in g84-vIP and 580 proteins in GS15-vIP **(Figure 2, Table S3)**. The GO analysis of all three vesicle IPs from control cells revealed that the vesicular proteins primarily belong to ER-Golgi and intra Golgi vesicle transport and glycosylation pathways **(Figure 2A-C, left panels, Table S3)**. In addition, an enrichment in “intermediate filament organization pathway” was detected in both g84-vIP and GS15-vIP vesicles, indicating potential involvement of intermediate filaments in the intra-Golgi vesicular trafficking. Analysis of the second vesicle IP dataset confirmed the primary involvement of immunoisolated proteins in Golgi trafficking and glycosylation **(Supplementary Figure 2A-C, left panels, Table S3)**. GO enrichment analysis for the cellular component revealed major enrichment in proteins associated with Golgi apparatus sub-compartments and Golgi membrane (**Figure 2A**-**C** **right panels, Supplementary Figure 2A**-C **right panels, Table S3),** validating our vesicle immunoprecipitation strategy. The GO enrichment analysis for biological processes and cellular components of CCD vesicles immunoisolated from COG4 depleted cells (G-CCDvIP, g84-CCDvIP, GS15-CCDvIP, and V7-CCDvIP) revealed similar results **(Figure 3A, B, C and Supplementary Figure 3A, B, C, Table S3)**. This suggests that acute COG inactivation did not mislocalize transmembrane vesicle “handles,” and all analyzed vesicle populations originated from Golgi membranes.

**Figure 2.**
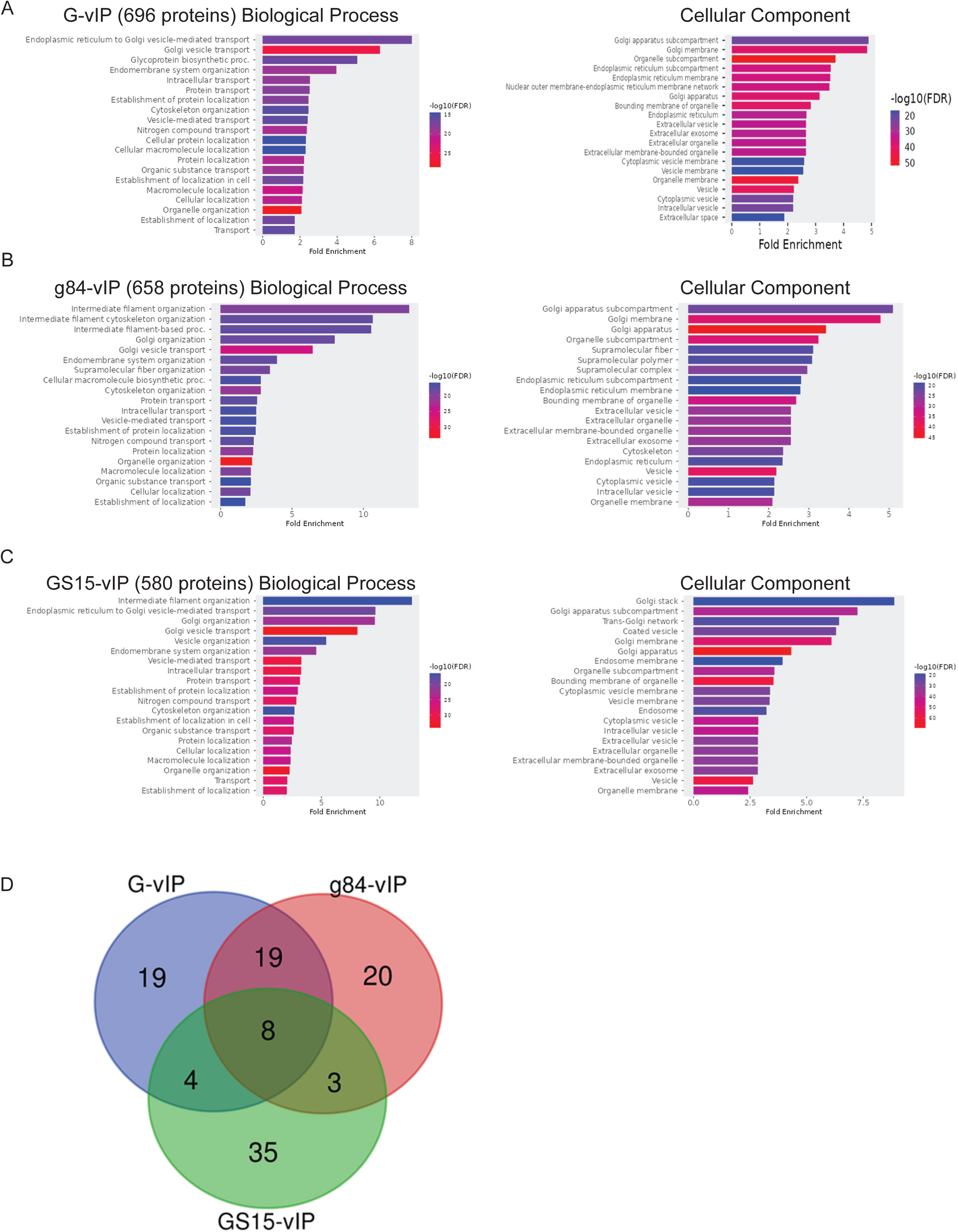
Composition of different populations of vesicle proteome isolated from control cells. **A)** Gene Ontology (GO) term enrichment analysis of biological process and cellular components for proteins enriched (≥2-fold, P ≤0.05) in **A)** Giantin pull-down vesicles (G-vIP), **B)** Golgin-84 pull-down vesicles (g84-vIP) and **C)** GS15 pull-down vesicles (GS15-vIP) isolated from control cell compared to input (S30). **D)** Venn diagram depicting the overlap of top 50 enriched transmembrane proteins in Giantin (G-vIP), Golgin-84 (g84-vIP), and GS15 pull-down vesicles (GS15-vIP) in steady state (control cell).

**Figure 3.**
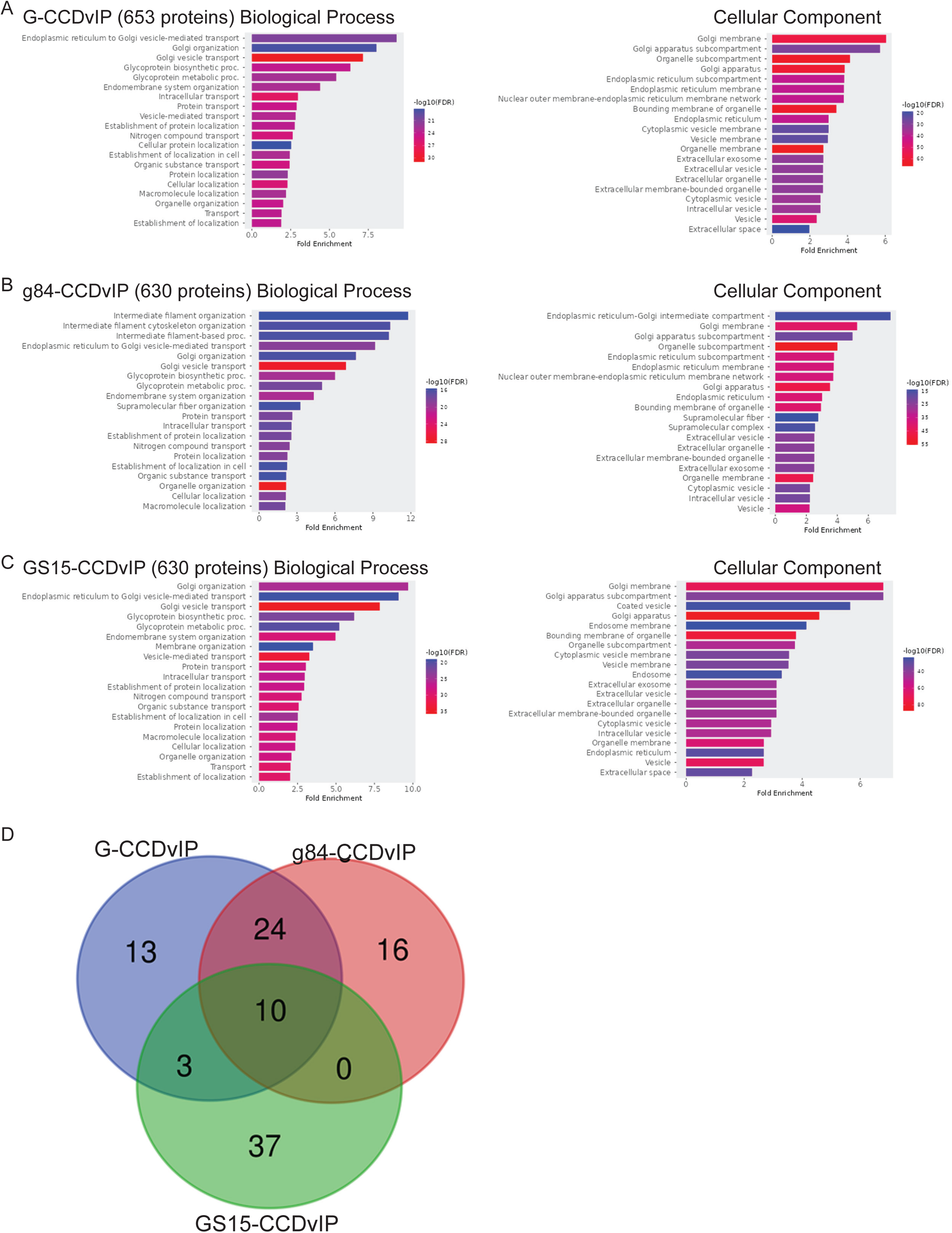
Composition of different populations of vesicle proteome isolated from Acute COG-depleted cells. **A)** Gene Ontology (GO) term enrichment analysis of biological process and cellular components for proteins enriched (≥2-fold, P ≤0.05) in **A)** Giantin pull-down vesicles (G-vIP), **B)** Golgin-84 pull-down vesicles (g84-vIP) and **C)** GS15 pull-down vesicles (GS15-vIP) isolated from acute COG4 depleted cell compared to input (S30). **D)** Venn diagram depicting the overlap of top 50 enriched transmembrane proteins in Giantin (G-CCDvIP), Golgin-84 (g84-CCDvIP), and GS15 pull-down vesicles (GS15-CCDvIP) in acute COG4 depleted cells.

### Characterization of immunoprecipitated vesicle populations

To identify the specific “molecular signatures” of the immunoprecipitated vesicle populations, we first analyzed the composition of the top 50 transmembrane proteins enriched in each vesicle-IP compared to the input (S30). We hypothesized that some peripheral proteins might be removed from the membrane surface due to multiple wash steps during vesicle isolation. In contrast, TM proteins, which are stably embedded in the vesicle membrane, would remain associated. This approach allowed us to focus on the most reliably enriched membrane-bound proteins in each vesicle population. In the first set of vesicle IP **(Table S3)**, we found that one TM protein, the glycosylation enzyme ST6GALNAC4, was detected only in vIPs, while others were enriched 15-100-fold. In all six vesicle groups isolated from control and COG4-depleted IPs, 64-84% of the most enriched TM were affiliated with the “Golgi set”, validating the vesicle isolation approach. The analysis revealed that the molecular signature of G-vIP, g84-vIP, and GS15-vIP vesicles isolated from control cells was significantly distinct, with only eight proteins being shared by all three vesicle populations **(Figure 2D)**. Common proteins include intra-Golgi v-SNARE GS28/GOSR1 (D’Souza et al., 2023; Sumya, Pokrovskaya, D’Souza, et al., 2023) and a putative Golgi autophagy receptor YIPF4 (Hickey et al., 2023). 32 Golgi proteins were among 50 most enriched TM proteins in the G-vIP fraction **(Table S3)**. This set contains 14 enzymes and sugar transporters, including cis-medial Golgi N-glycosylation enzymes MAN1A2, MGAT2, FUT8, O-glycosyltransferases GALNT2, 7, 10, and a 3’-phosphoadenylyl sulfate:adenosine 3’,5’-bisphosphate antiporter SLC35B2.

g84-vIP membranes were enriched with 42 Golgi TM proteins, including 18 enzymes and sugar transporters from both N-glycosylation (MAN2A1, MGAT2) and O-glycosylation (GALNT2, B4GAT1) pathways **(Table S3)**. The presence of cis (MAN1A2), medial (MGAT1), and trans-Golgi (CHST14) resident enzymes indicates that g84-vIP trafficking intermediates are employed for protein recycling from multiple Golgi compartments.

GS15-vIP membranes were enriched in 42 Golgi resident TM proteins, including several trans-Golgi enzymes and sugar transporters (B4GALT1, B4GALT5, BPNT2, EXTL3, SLC35E1) as well as TGN SNAREs STX16 and VTA1A, cargo receptors TGOLN2/TGN46, MP6R, IGF2R, and SORT1 and autophagy protein ATG9A **(Table S3)**. The molecular signature of GS15-vIP vesicles indicate that these carriers are preferentially derived from TGN/trans-Golgi. The presence of multiple species of Golgi enzymes in different vesicle populations isolated from control cells supports the model of continuous vesicular recycling of Golgi resident proteins.

The second round of vesicle IP employed a more sensitive mass-spec instrument and detection program, resulting in a higher protein detection rate, with many proteins identified exclusively in the vesicle population. Consequently, to identify molecular signatures of immunoprecipitated vesicles, we conducted an analysis of the 100 most enriched TM proteins. A comparison of Giantin, VAMP7, and GS15 vIPs revealed that only 24% of TM proteins were common in all three groups, indicating that they represent significantly distinct vesicle populations **(Supplementary Figure 2D, Table S3)**. Common proteins include SNAREs (SEC22A, VAMP4), Golgi enzymes (B4GALT7, GALNT11), and transporters (SLC9A7, SLC30A6, SLC35B1, and SLC35E1) **(Table S3)**.

77 Golgi proteins were among the 100 most enriched TM proteins in the G-vIP fraction. This set contains 28 enzymes and sugar transporters, including cis-medial Golgi N-glycosylation enzymes MAN1A2, MANEA, MGAT4B, O-glycosyltransferases GALNT4, 7, 11, 13, and nucleotide sugar transporters SLC35A5, SLC35B1.

The V7-vIP (VAMP7 vesicles) set was enriched for 84 Golgi TM proteins, including 18 enzymes and sugar transporters, mainly from the O-glycosylation pathway (GALNT4, GALNT7, GALNT11, B3GNT9). GS15-vIP was enriched for 62 Golgi resident TM proteins, including 18 trans-Golgi enzymes and sugar transporters (ST3GAL1, CHST3, ST8SIA6, SLC35A5) as well as trans-Golgi/TGN SNAREs STX10, STX16, VAMP4 and VTA1A, and cargo receptors TGOLN2/TGN46, MP6R **(Table S3)**.

After acute COG4 depletion, the molecular signature of each vesicle population was slightly altered with an increase in Golgi resident proteins and, most notably, components of the glycosylation machinery **(Figure 3D, Table S3)**. Comparison of the molecular signatures for G-CCDvIP, g84-CCDvIP and GS15-CCDvIP showed an increase from 8 to 21 of common proteins. Similarly, in the second vesicle IP, the comparison of vesicle protein content revealed that the overlap between G-CCDvIP, V7-CCDvIP, and GS15-CCDvIP increased from 21 shared proteins compared to control vIP to 42 in the CCD vesicles **(Supplementary Figure 3D, Table S3)**.

### COG depletion leads to a quantitative increase in the vesicle-associated glycosylation machinery

Since the molecular signatures of the three isolated Golgi vesicle pools were significantly distinct, we proceeded to compare the total proteomes of each CCD-vIP with their corresponding control vesicles. This comparison aimed to identify those Golgi proteins that were specifically enriched or depleted in the COG-depleted vesicles, providing insights into how COG complex dysfunction affects the composition and function of intra-Golgi vesicular trafficking intermediates. This analysis allowed us to determine the extent by which the Golgi resident proteins and trafficking machinery depend on COG function for proper recycling and localization. Notably, 87% of the glycosylation machinery was significantly increased in G-CCDvIP vesicles **(Figure 4B, Table S4)**. Among glycosyltransferases, the highest increase was detected for medial-Golgi enzymes CHST11 (22-fold), MGAT5 (18-fold), and MAN1A1 (14-fold) **(Table S4)**. The volcano plot illustrates the accumulation of GALNT2, MGAT2, MAN1A1, and MAN2A1 enzymes in G-CCDvIP **(Figure 5A, Table S1, S4)**. WB analysis of Giantin pull-down vesicles revealed a significant increase of MGAT1 and GALNT2 and a moderate increase of B4GALT1 **(Figure 5B)**, corroborating the mass spectrometry data. Conversely, only 39% of the other Golgi transmembrane proteins showed an increase in G-CCDvIP **(Figure 4A, 4C)**. The most substantial increases were observed for Glutaminyl-peptide cyclotransferase-like protein QPCTL (11-fold) and soluble calcium-activated nucleotidase CANT1 (10-fold) (**Table S4)**. Several TM proteins, including CAV1, GJA1, LDLR, SURF4, and VAMP2, exhibited decreased abundance in G-CCDvIP vesicles, suggesting that COG does not regulate their trafficking. Furthermore, although only 28% of the luminal Golgi residents accumulated in G-CCDvIP **(Figure 4D)**, several, including FAM3A, STC2, DIPK2A, and SDF4, showed a 7-10-fold increase in abundance. The majority (80%) of peripheral membrane proteins did not show significant changes in abundance in G-CCDvIP, with the exception of the vesicular coat components ARF1, ARF3, AP1B1, and COPG2. These coat proteins exhibited a marked decrease in abundance, suggesting that the disassembly of vesicle coats from the surface of stalled vesicles **(Figure 4E, Table S4)**. Among the 21 Golgi Rab proteins detected in G-CCDvIP, only 5 Rabs (Rab2A, Rab2B, Rab7A, Rab11B, and Rab15) showed some level of increase, indicating that the majority of Rab proteins did not recycle in Giantin-positive vesicles in a COG-dependent manner.

**Figure 4.**
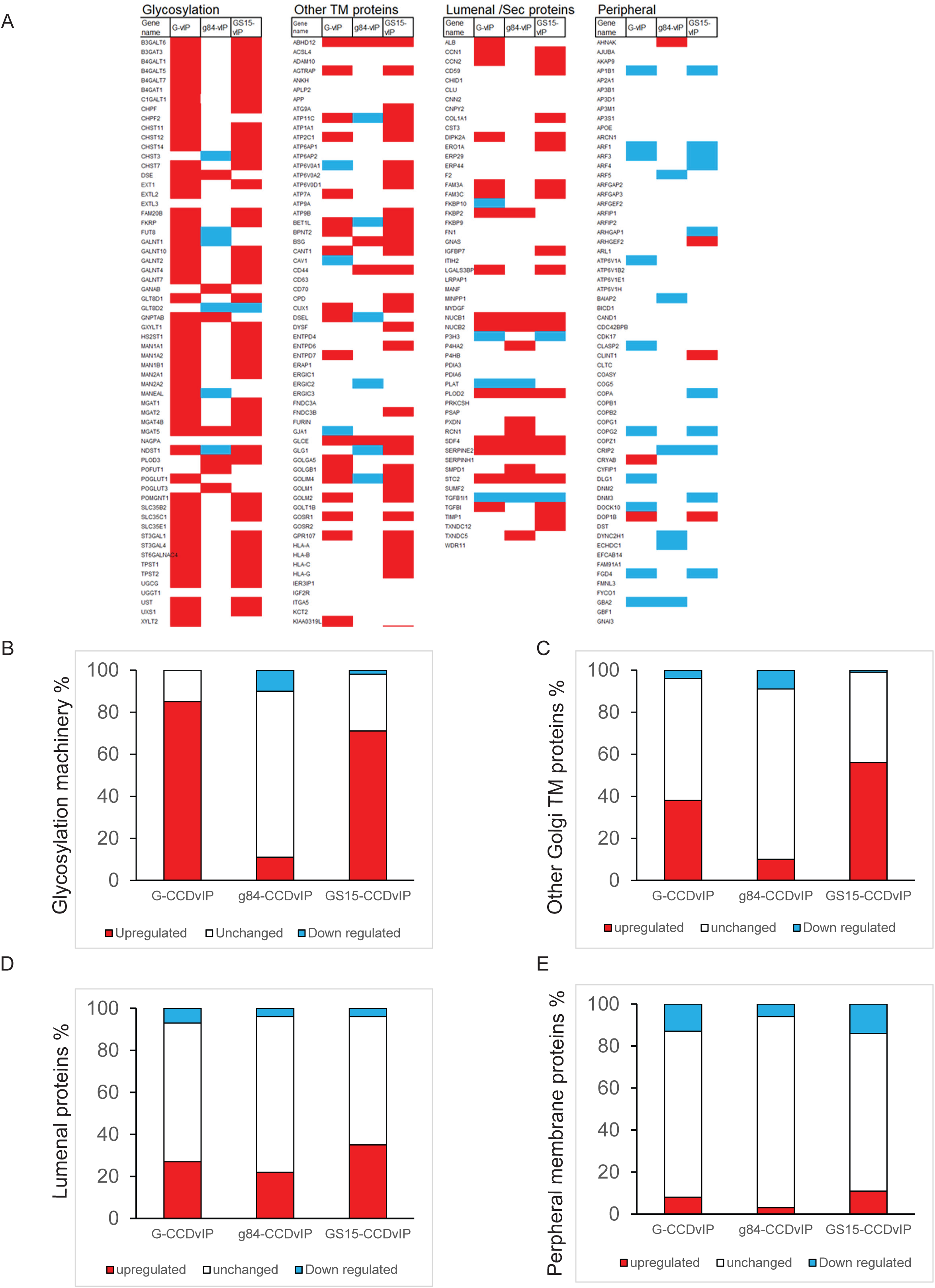
The CCD vesicles comprise different types of Golgi resident proteins. **A)** The captured image from >Table 2 indicates the upregulation (Red), downregulation (Blue), and unchanged portion (transparent) of Golgi resident proteins in CCD vesicles pull-down by Giantin (G-CCDvIP), Golgin-84 (g84-CCDvIP), and GS15 (GS15-CCDvIP) compared to control. The bar graph shows the percentage of upregulated, unchanged, and downregulated **B)** Glycosylation machinery, **C)** Other transmembrane proteins **D)** Lumenal secretory proteins, and **E)** Peripheral membrane proteins in G-CCDvIP, g84-CCDvIP, and GS15-CCDvIP compared to G-vIP, g84-vIP, and GS15-vIP, respectively.

**Figure 5.**
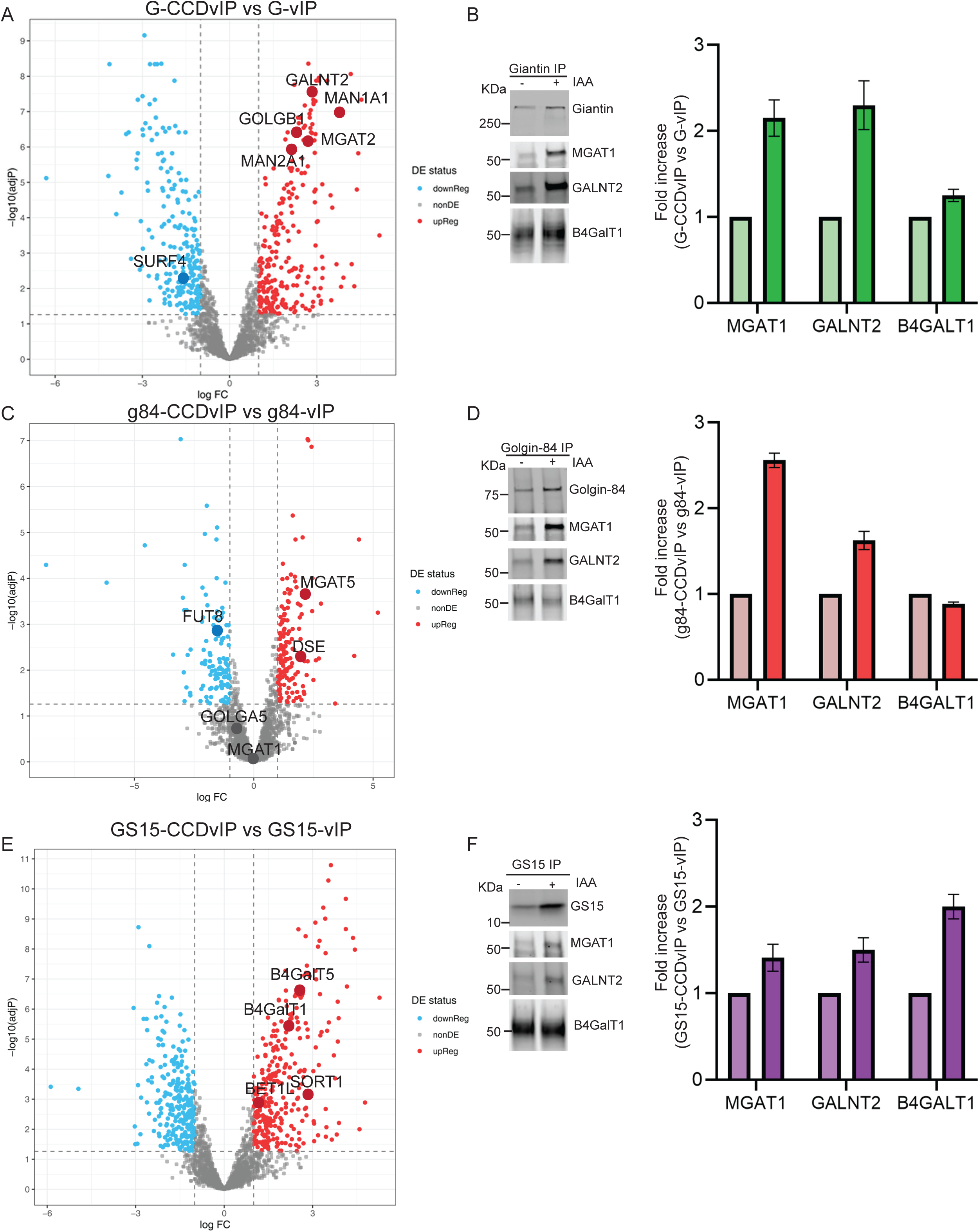
Characterization of the different populations of vesicle proteome isolated from control and acute COG4-depleted cells. Label-free mass spectrometry analysis of G-vIP/G-CCDvIP, g84-vIP/g84-CCDvIP, and GS15-vIP/GS15-CCDvIP respectively. **A)** Volcano plot depicting the fold changes of proteins from four independent experiments of a direct comparison of G-vIP (control cells) and G-CCDvIP (acute COG4 depleted cells). **B)** The left panel shows the WB analysis of Giantin, MGAT1, GALNT2, and B4GALT1 in Giantin pull-down vesicles before and after IAA treatment. The right panel shows the quantification of fold enrichment of MGAT1, GALNT2, and B4GALT1 in G-CCDvIP compared to G-vIP. **C)** Volcano plot depicting results from four independent experiments of a direct comparison of g84-vIP (control cells) and g84-CCDvIP (acute COG4 depleted cells). **D)** The left panel shows the WB analysis of Golgin-84, MGAT1, GALNT2, and B4GALT1 in Golgin-84 pull-down vesicles before and after IAA treatment. The right panel shows the quantification of fold enrichment of MGAT1, GALNT2, and B4GALT1 in g84-CCDvIP compared to g84-vIP. **E)** Volcano plot depicting results from four independent experiments of a direct comparison of GS15-vIP (control cells) and GS15-CCDvIP (acute COG4 depleted cells). **F)** WB analysis of GS15, MGAT1, GALNT2, and B4GALT1 in GS15 pull-down vesicles before and after IAA treatment. The upper right panel shows the quantification of MGAT1, GALNT2, and B4GALT1 fold enrichment in GS15-CCDvIP compared to GS15-vIP. In volcano plots, data are from four replicates each. Colors indicate red, fold-change >2 and p <0.05; grey, fold-change >2 or <2 and p <0.05; blue, fold-change <2 and p <0.05.

Notably, COG4 depletion did not result in a significant accumulation of Golgi resident proteins in g84-CCDvIP vesicles **(Figure 4A, Table S4)**. Among glycosyltransferases, the most significant increase in abundance was detected for MGAT5 (5 fold) and DSE (4 fold), however 89% of glycosylation machinery did not change **(Figure 4C, Table S4)** or even decreased (FUT8, GALNT1, CHST3, NDST1, MANEAL) in abundance, indicating that golgin84-positive membrane carriers are not widely utilized for COG-dependent recycling **(Figure 5C, Table S1, S4)**. WB analysis of golgin84 pull-down vesicles showed an increase in MGAT1 and GALNT2 **(Figure 5D)**. The lack of changes in this vesicle pool is puzzling but could be related to a significant accumulation of golgin-84 protein in G-CCDvIP vesicles.

An analysis of GS15-CCDvIP revealed that a substantial fraction (69%) of the glycosylation machinery increased in abundance in GS15-CCDvIPs **(Figure 4B, Table S4)**. A significant increase in abundance was detected for medial-trans Golgi enzymes MAN1A1 (22-fold), NDST1 (17-fold), and CHST12 (15-fold) (**Table S4)**, indicating that COG dysfunction may cause the rerouting of early enzymes to later Golgi compartments. In the volcano plot, we observed the accumulation of B4GALT1, B4GALT5, and SORT1 in GS15-CCDvIP compared to GS15-vIP **(Figure 5E, Table S1, and S4)**. WB analysis of GS15 pull-down vesicles indicated an observable increase in B4GALT1 and a moderate increase in MGAT1 and GALNT2 enzymes in GS15-CCDvIP **(Figure 5F)**. In addition to the glycosylation machinery, 56% of other TM proteins were also increased in abundance in GS15-CCDvIP **(Figure 4C, Table S4)**. The most notable increase was observed for QPCTL (39-fold), SFT2D3 (12-fold) and BSG (12-fold) **(Figure 4A, Table S4)**. More than a third (35%) of luminal Golgi resident proteins increased in abundance in GS15-CCDvIPs **(Figure 4D, Table S4)**, indicating that GS15-positive transport vesicles, along with Giantin-positive, serve as primary retrograde transport carriers in the COG-dependent intra-Golgi recycling pathway. There was no increase in peripheral membrane proteins in GS15-CCDvIP **(Figure 4E)**.

Several Golgi TM proteins did not show any change (SLC30A7, UGGT1, ATP9A, CD63, ERGIC1, ERGIC3, FURIN, SEC22B), or even decreased significantly in abundance (GLT8D2, ERGIC2, LMAN1, SURF4) in CCDvIPs, indicating that these proteins do not recycle in COG-dependent trafficking intermediates.

Analysis of the second round of vesicle precipitations revealed a similar trend with an increase in Golgi resident proteins **(Supplementary Figure 4B, Table S4)**. The results revealed that 86% of the glycosylation machinery proteins was increased in abundance in all three vesicle pools (G-CCDvIP, V7-CCDvIP, and GS15-CCDvIP) compared to the control **(Supplementary Figure 4A, 4B, Table S4)**, agreeing with the results obtained by the first vesicle isolation. The abundance of Golgi TM proteins unrelated to glycosylation showed little change in all CCD-vIPs, suggesting that the COG-dependent vesicles are not involved in their recycling **(Figure Supplementary 4A, 4C, Table S4)**. The majority of luminal and peripheral membrane proteins also did not show any significant changes in CCD-vIPs, with the notable exception of coat components such as AP1S1, ARF1, ARFGAP1, and all COPI subunits, which showed a marked decrease in abundance. **(Supplementary Figure 4A, 4D, 4E)**. This finding corroborates the results from the first vesicle-IP, confirming that coat disassembly is a specific response to COG complex dysfunction.

Notably, several components of the glycosylation machinery (CHST10, PXYLP1, and ST6GALNAC3) were detected only in the G-CCDvIP and not in the G-vIPs, indicating exceptional enrichment of these proteins in vesicles upon COG depletion. Over 50 other enzymes and sugar transporters were significantly enriched in G-CCDvIPs compared to Giantin-IP vesicles isolated from control cells **(Table S4)**. The volcano plot revealed significant accumulation of cis-medial enzymes in G-CCDvIPs **(Supplementary Figure 5A)**. Western blot analysis showed a substantial increase in GALNT2 and a moderate increase in MGAT1 and B4GALT1 in G-CCDvIP **(Supplementary Figure 5B)**.

The composition of V7-vIP trafficking intermediates also experienced a significant increase (>50%) in the enrichment of components of Golgi glycosylation machinery **(Supplementary Figure 4B)**. At least 26 members of the glycosylation machinery from various Golgi cisternae were detected in V7-vIP only after COG4 depletion, making it impossible to quantify the extent of their enrichment. Notable increases were observed for C1GALT1C1 (16-fold), C1GALT1 (13-fold), and POMGNT1 (11-fold) **(Table S4)**. The volcano plot of V7-CCDvIP vs V7-vIP showed accumulation of GALNT7, MGAT1, GALNT4, B3GANT9 **(Supplementary Figure 5C)**. Western blot analysis indicated an observable increase in GALNT2 and a moderate increase in B4GALT1 enzymes in V7-CCDvIP **(Supplementary Figure 5D)**.

The Golgi proteome of GS15-CCDvIP showed moderate changes like those observed in the first experiment **(Supplementary Figure 5E, 5F)** highlighted by the increase in members of Golgi glycosylation machinery **(Supplementary Figure 4B),** particularly enzymes localized in trans-Golgi compartments **(Table S4)**. Enrichment in trans-Golgi enzymes (B4GALT1, CHST14), SNAREs, and mannose-6-receptors M6PR and IGF2R supports the notion that GS15-CCDvIP trafficking intermediates preferentially originate from late Golgi sub compartments.

### Preferential accumulation of Golgi Glycosylation machinery in CCD vesicles

Comparison of proteomes from the G-CCDvIP, g84-CCDvIP, and GS15-CCDvIP vesicle populations reveal that most of the Golgi resident proteins tend to accumulate in G-CCDvIP and GS15-CCDvIP. In g84-CCDvIP, only golgin84/GOLGA5 showed specific accumulation compared to the other two vesicle populations. The majority of Golgi residents which accumulated in the G-CCDvIP consisted of cis-medial Golgi enzymes, including MGAT1, MGAT2, MAGT4b, MAN1A2, MAN2A1, GALNT1, GALNT2, GALNT7, and GALNT10 **(Figure 4A, Table S4)**. The largest set of Golgi proteins (63) were specific to GS15-CCDvIP vesicles, and included several trans-Golgi enzymes (B4GALT1, B4GALT5, ST3GAL4, CHST3, CHST11, CHST12, NAGPA) and a significant number of Golgi SNARE proteins **(Figure 4A, Table S4)**. A side by side comparison of protein enrichment in the different populations of CCD vesicles from the second vesicle IP set was generally in agreement with the results from the first Golgi vesicle pull-down. A comparison of the protein composition of G-CCDvIP with GS15-CCDvIP vesicles confirmed that cis-medial localized components of the glycosylation machinery preferentially accumulate in Giantin-positive transport carriers. Similar to the first vesicle IP, MAN1A2, MGAT2, MGAT4B, GALNT2, GALNT7 and GALNT10 were enriched in G-CCDvIP **(Supplementary Figure 4A, Table S4)**. V7-CCDvIP did not exhibit specific enrichment in Golgi recycling proteins when compared to G-CCDvIP **(Supplementary Figure 4, Table S4**), suggesting a significant overlap between the two and indicating that V7-CCDvIP likely represents a sub-population of G-CCDvIP intermediates.), suggesting a significant overlap between the two and indicating that V7-CCDvIP likely represents a sub-population of G-CCDvIP intermediates.

### Non-tethered CCD vesicles specifically captured by ectopically expressed golgin-84

To test the functionality of CCD vesicles, we employed their capture by golgins relocated to mitochondria. The Munro lab has reported that several golgins, including golgin-84, when relocated to mitochondria, can capture intra-Golgi vesicles (Wong & Munro, 2014). This capture required disruption of the Golgi ribbon using the microtubule depolymerization agent nocodazole, likely to facilitate the close spatial association between mitochondria covered with golgin-84 and vesicles budding from multiple Golgi ministacks. We hypothesized that acute COG inactivation increases the number of non-tethered vesicles, thereby removing the need for disruption of the Golgi structure. Indeed, golgin-84-decorated mitochondria effectively captured CCD vesicles carrying Giantin and GALNT2 (**Fig.6A**). This efficient vesicle capture required COG4 depletion (**Fig 6A, 6B**) and was specific; vesicles carrying GS15 and B4GalT1 were not captured by ectopically expressed golgin-84 (**Figure 6D**). Furthermore, an expression construct lacking the golgin-84 module did not capture any CCD vesicles (**Fig 6C**.). Our data confirm the ability of golgin-84 to capture a specific subset of intra-Golgi vesicles identified by the Munro lab and clearly indicates that CCD vesicles that transiently accumulate in COG-deficient cells are identical to the transport intermediates present in wild-type cells. As expected, expressing golgin-84 on the outer mitochondrial membrane in COG4-depleted cells “glued” vesicles between mitochondria, resulting in an aggregation/fragmentation phenotype. A similar phenotype has previously been observed by us and others in HeLa cells expressing either COG subunits (Willett, Kudlyk, et al., 2013) or golgins (Wong & Munro, 2014) on the mitochondrial surface.

**Figure 6.**
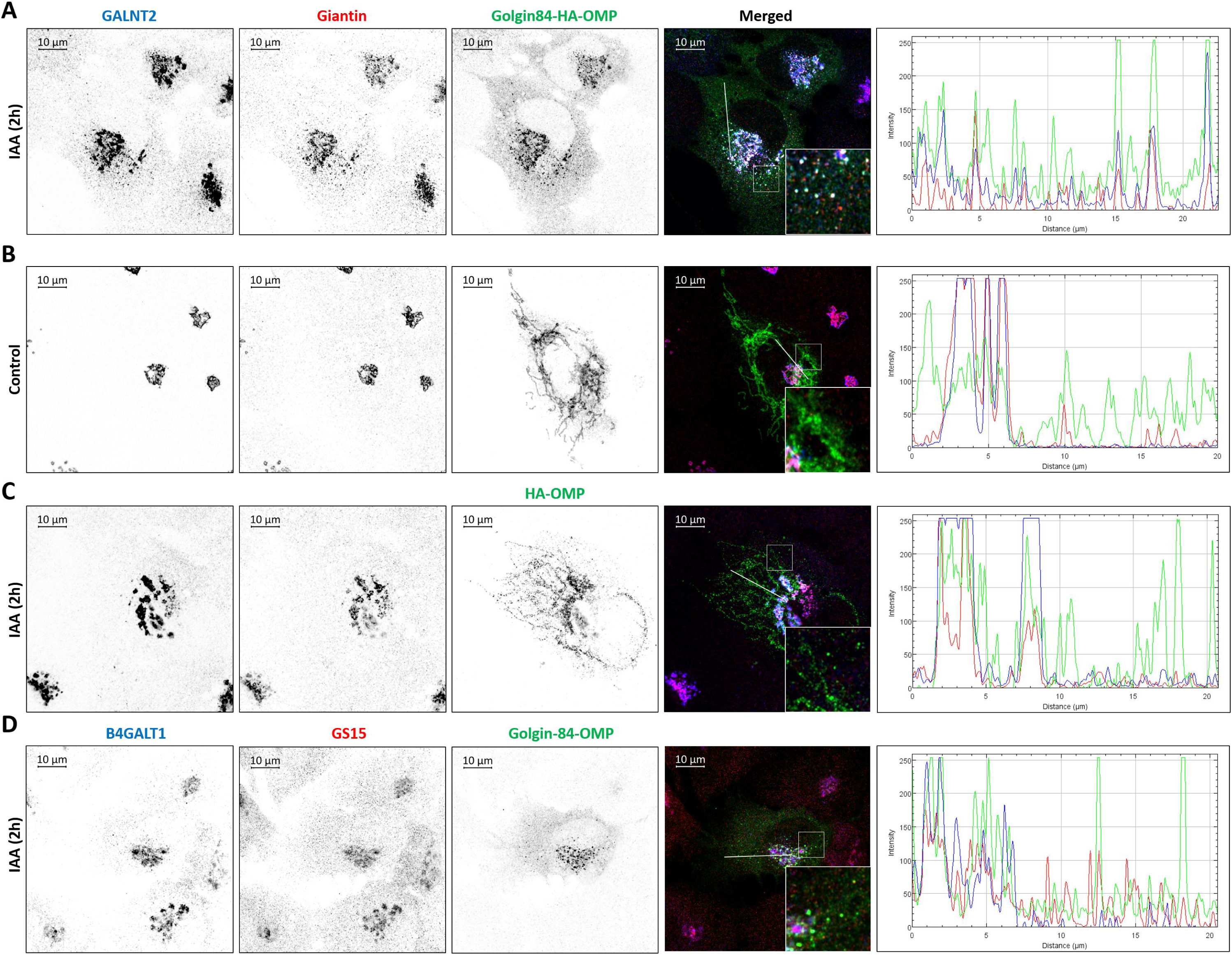
CCD vesicles are functional transport intermediates that can be rerouted to golgin-decorated mitochondria. Airyscan superresolution immunofluorescence (IF) analysis of RPE1 COG4-mAID cells transiently expressing either golgin84-HA-OMP25 or HA-OMP25-HA. Scale bars represent 10μm. Far right panels are the line scan plots of relative intensity of the designated distance. Prior to fixation, cells expressing golgin84-HA-OMP25 were either **A)** treated with IAA for 2h (depleted COG4) or **B)** untreated (intact COG4/Control). Cells were then stained for the golgin84-capturable vesicle components GALNT2 (Blue) and Giantin (Red). Cells were also stained for HA (Green) to visualize the ectopically expressed golgin-84. **C)** Cells expressing HA-OMP25 were treated with IAA for 2h prior to fixation and then stained for GALNT2 (Blue), Giantin (Red), and HA (Green). **D)** Cells ectopically expressing golgin-84 were treated with IAA for 2h prior to fixation and then stained for vesicle components not known to be captured by golgin-84, B4GALT1 (Blue) and GS15 (Red). Cells were also stained for HA (Green).

## Discussion

In this study, we performed an unbiased analysis of the Golgi-derived vesicle proteome by immunoisolating endogenous trafficking intermediates and conducting high-resolution data-independent acquisition mass spectrometry. This approach identified 776 proteins from a curated “Golgi set” (Fasimoye et al., 2023), with 386 Golgi proteins being significantly enriched in the isolated vesicles. The Golgi proteome coverage was comparable to that of the recently published Golgi-IP method (Fasimoye et al., 2023), demonstrating high efficiency of the vesicle-IP approach. By utilizing three distinct vesicle markers from different Golgi sub-compartments, we separated the vesicle pool into three semi-independent populations of Golgi trafficking intermediates, permitting a detailed analysis of intra-Golgi recycling. Additionally, acute depletion of the COG vesicles tethering machinery enabled us to focus on trafficking intermediates and Golgi resident proteins that depend on the COG complex for recycling, localization, and function. Our findings suggest that nearly all components of the Golgi glycosylation machinery are COG-dependent for proper recycling and localization.

Ontology analysis of the proteomes from all twelve vesicle-IP experiments revealed a significant enrichment of proteins involved in ER-Golgi and intra-Golgi vesicle transport and glycosylation processes. Despite this common trend, there were notable differences in the protein composition of G-vIP, g84-vIP, and GS15-vIP. While most Golgi resident proteins were detected across all three vesicle populations, cis and medial TM proteins were predominantly found in G-vIP, whereas trans-Golgi TM proteins were more abundant in GS15-vIP. The third vesicle pool, identified as g84-vIP in the first vesicle MS and VAMP7-vIP in the second, was distinct from both G-vIP and GS15-vIP, indicating that intra-Golgi trafficking is mediated by at least three different vesicle carrier types.

Overlap in vesicle molecular signatures increased upon COG depletion, suggesting that persistent defects in vesicle tethering compromise intra-Golgi sorting mechanisms. Furthermore, the presence of nearly all species of Golgi enzymes across the different vesicle populations in wild-type cells supports the concept of continuous vesicular recycling of the Golgi glycosylation machinery. More than 92% of Golgi glycosylation enzymes, and 68% of other TM proteins, showed a significant increase in abundance in at least one type of CCD-vIP, indicating that the recycling of these proteins is dependent on COG function. Given that COG-KO cells lose more than 90% of Golgi enzymes (Bailey Blackburn et al., 2016; Sumya, Pokrovskaya, D’Souza, et al., 2023), we conclude that the entire Golgi glycosylation machinery relies on COG-dependent vesicular recycling for its localization and function.

Approximately 48% of lumenal proteins showed a significant increase in at least one type of transport vesicle, indicating that nearly half of Golgi lumenal proteins are recycled via CCD vesicles. Notably, Golgi resident proteins NUCB1/2 and Cab45/SDF4 were significantly increased in CCD vesicles, whereas the levels of transiently passing cargo, such as secretory proteins prosaposin (PSAP) and SERPINEH1, remained relatively unchanged.

Since the role of intra-Golgi vesicular trafficking in delivering secretory proteins remains unclear, we compared the proteins accumulated in CCD vesicles with those previously identified in the RPE1 secretome (Sumya et al., 2021). To eliminate potential contaminants, we manually selected soluble secretory proteins with known signal sequences using data from the UniProt database (Apweiler et al., 2004). A comparative analysis of the resulting 360 proteins showed that 11% and 18% of “soluble secretory proteins” were significantly enriched in CCD vesicles in experiments 1 and 2, respectively (**Supplementary Figure 6, Table S5**).

Interestingly, only 20 proteins were common between both experiments, including ten known Golgi-resident lumenal proteins. Furthermore, only nine proteins (CFB, DCD, LAMB1, LOX, NT5E, PLOD3, PTX3, SERPINE1, and THBS1) that are not typically found in intracellular compartments were significantly increased in CCD vesicles. One possible explanation for the presence of specific secretory proteins in CCD vesicles is their prolonged interaction with the glycosylation machinery due to the extensive processing required for glycosylation. This strongly suggests that CCD vesicles primarily recycle Golgi-resident proteins rather than proteins destined for secretion.

Analysis of peripheral membrane proteins in the vesicle proteome revealed the presence of at least three vesicular coats—COPI, AP1, and AP3—indicating that the immunoisolated vesicles were formed through multiple sorting/budding mechanisms. It has been previously proposed that transport vesicles remain coated until the coat is recognized by a specific tether (Cai et al., 2007). Interestingly, several coat components, including ARF1, COPA, COPG2, and AP1B1, were significantly reduced in vesicles isolated from COG-deficient cells, implying that stalled vesicles lose their coats in the absence of coat-tether interaction. These findings suggest that the vesicle-forming machinery initially bound to recycling vesicles becomes dissociated from trafficking carriers which are unable to tether and fuse with Golgi cisternae.

Since the COG complex interacts with the COPI coat (Oka et al., 2004; Suvorova et al., 2002; Willett et al., 2014) and proteomics data suggest that at least some CCD intermediates are formed by COPI machinery, we compared the vesicle proteomics data obtained in this study with the proteome of in vitro formed COPI vesicles (Adolf et al., 2019). Comparison of the 277 Golgi proteins enriched in CCDvIP **(Table S6)** with the 207 proteins identified in COPI vesicles isolated from HeLa cells revealed that 109 Golgi proteins are common to both **(Supplementary Figure 7A)**. The remaining 166 proteins are unique to CCD-vIP, suggesting that these proteins are incorporated into COPI vesicles only under in vivo conditions or are recycled in trafficking intermediates formed by AP coats. Further comparison of the CCD-vIP proteome with the 38 Golgi proteins common to all COPI vesicle proteomes from HeLa, HepG2, and iMΦ cell lines, revealed that more than 90% of COPI vesicle core proteome is shared with CCD-vIP **(Supplementary Figure 7B).** The remaining three proteins unique to the COPI vesicle proteome—KDELR1, SLC30A6, and ERP44 were detected in vIPs, but their abundance did not increase in COG4-depleted cells, indicating their independence from the COG complex machinery. KDELR1 and ERP44 are known to recycle through the ER, likely via the DSL1/NZR-dependent pathway (Lewis & Pelham, 1992; Tempio & Anelli, 2020). The Zn2+ transporter ZnT6/SLC30A6, which regulates ERP44 activity (Amagai et al., 2023), likely follows the same COG-independent recycling route. Thus we conclude that a significant fraction of the CCD-vIP Golgi proteins recycle in a vesicle formed by COPI coat machinery.

Our findings support a model in which the Golgi residents recycle in separate, and distinct vesicles formed from the different cisternae through various sorting and budding mechanisms. The entire Golgi glycosylation machinery, along with a significant fraction of other Golgi residents, recycles in a COG-dependent manner, while a subset recycles independently of COG **(Figure 7).**

**Figure 7.**
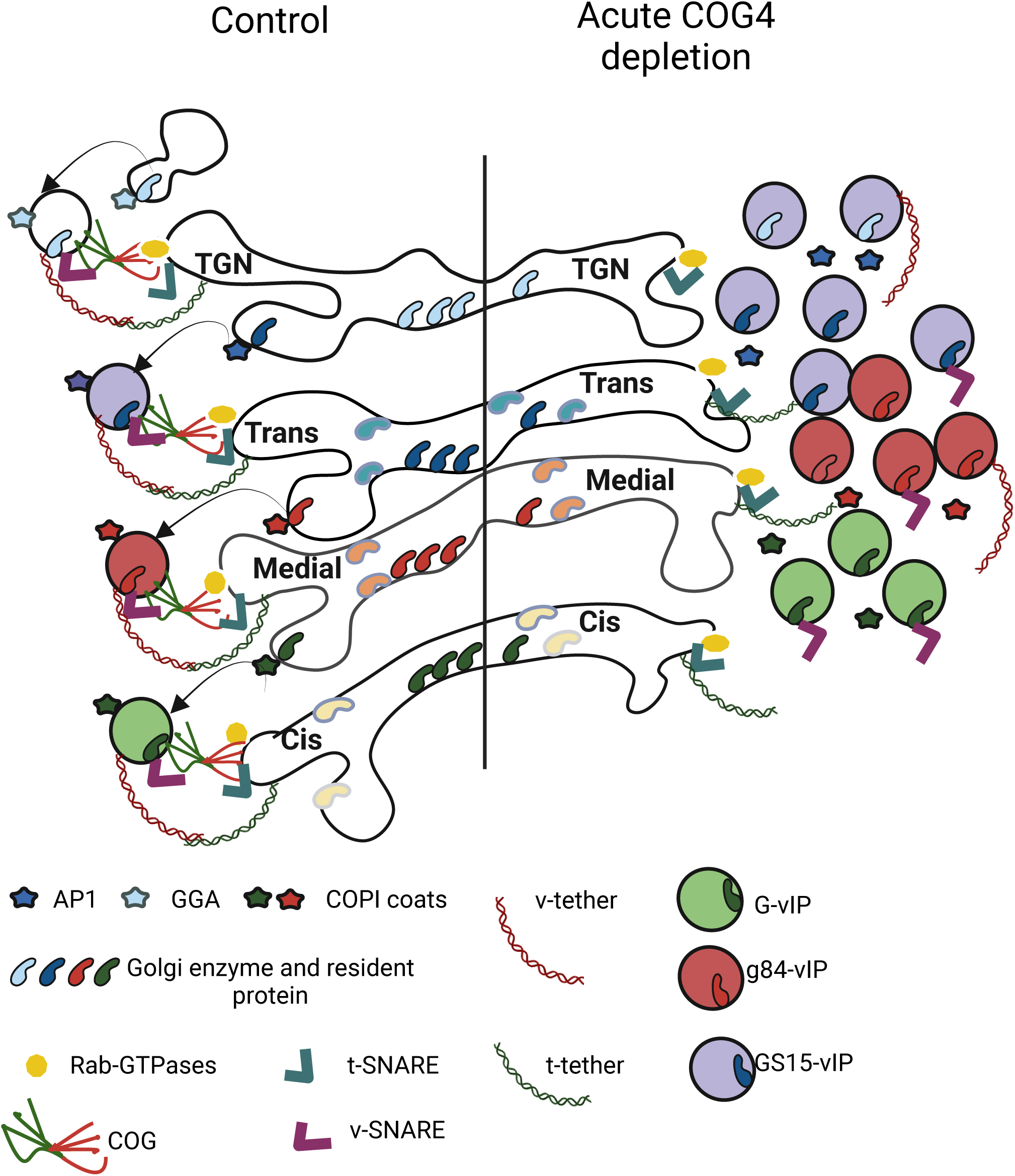
Model depicting the intra-Golgi intermediates in control and Acute COG depleted cells. Some Golgi residents recycle in the COG-dependent vesicles (CCD vesicles), while others recycle independently as they remain in the Golgi after COG4 depletion.

The presence of the same Golgi resident protein in multiple vIP populations suggests that their intra-Golgi recycling itinerary involves several consecutive (from TGN to trans, from trans to medial, etc.) or complementary (from TGN, trans, and medial to cis) trafficking steps.

Previous studies have shown that mitochondria-relocated golgins can capture specific sets of trafficking intermediates (Wong & Munro, 2014). Notably, disruption of Golgi structure through nocodazole treatment was essential for capturing intra-Golgi vesicles by relocated golgins. In contrast, COG4 depletion accumulates non-tethered vesicles which can be captured by golgin-decorated mitochondria without the need for the nocodazole-induced Golgi disruption. The demonstration that giantin- and GALNT2-positive CCD vesicles are captured by ectopically relocated golgin-84 causing mitochondria aggregation, supports the overall conclusion that CCD vesicles are functional, freely diffusible trafficking intermediates. The mitochondrial aggregation test can be further developed by identifying additional components of the vesicle tethering machinery in cells where COG function has been acutely depleted.

To clarify the overlap between COG-dependent vesicles and vesicles captured by individual golgins, we compared the intra-Golgi vesicle proteome identified in this study with the proteome of vesicles captured by Golgin-97/GOLGA1 (Shin et al., 2020). A comparison of the 865 proteins enriched in CCDvIP (**Table S6**) with the 141 proteins identified in Golgin-97-tethered vesicles (G97-mito) revealed 33 shared proteins (**Supplementary Figure 7C**). Further analysis comparing the proteins enriched in GS15-CCDvIP (**Table S6**) with those in G97-mito captured vesicles showed that 32 proteins are common to both G97-mito and GS15-CCDvIP (**Supplementary Figure 7D**). This suggests that Golgin-97 may capture a subset of GS15-CCDvIP vesicles or that certain TGN proteins recycle through both intra-Golgi and endosome-to-Golgi pathways.

How does the COG complex regulate the tethering of such a diverse population of vesicles? COG is localized throughout the Golgi stack (Vasile et al., 2006) and interacts with multiple SNAREs and Rabs (Blackburn et al., 2019). Additionally, proximity biotinylation assays using endogenously expressed COG-TurboID fusions suggests that COG is in close proximity to nearly all coiled-coil vesicle tethers, golgins, which are localized on distinct Golgi cisternae (F.S and V.L unpublished data).

We propose a combinatorial vesicle tethering model in which the COG complex uses its C-terminal extending “arms” (Ha et al., 2016) to control the assembly of specific SNARE-Rab-golgin docking and fusion stations at the rim of each Golgi cisterna. In support of this model, we have recently identified two additional Golgi SNARE complexes: STX5/SNAP29/VAMP7 and STX5/VTI1B/STX8/YKT6 (D’Souza et al., 2023). In this model, deficiency in an individual v-SNARE, Rab, or golgin would compromise only a subset of intra-Golgi trafficking, and this defect could potentially be repaired by utilizing the remaining complementary components or alternative trafficking routes. Similarly, mutations in individual COG “arms” likely lead to partial effects. In contrast, dysfunction of the entire COG complex compromises multiple trafficking mechanisms, causing unreparable situations that result in missorting and degradation of the entire glycosylation machinery.

What is the machinery for recycling and localization of COG-independent proteins? One possibility is that they recycle in transport intermediates that are uniquely tethered by golgins. Indeed, STX5 interacts with p115/USO1 (Diao et al., 2008; Xu et al., 2002) and TGN46 vesicles can be captured by mitochondria-localized GOLGA1 (Wong & Munro, 2014). Another possibility is that COG-independent proteins are recycled via transient tubular connection between cisternae by a kiss-and-run mechanism (Mironov et al., 2013). The third possibility is that the itinerary of COG-independent proteins includes compartments outside the Golgi apparatus. For instance, STX5 was shown to recycle via ER (Hui et al., 1997), while TGN46 recycle via endosomal system (Banting & Ponnambalam, 1997). In this case their trafficking and localization should be controlled by ER-localized DSL1/NRZ and TGN-localized GARP complexes. In support of this scenario, TGN46 expression depends on GARP complex (Khakurel et al., 2024).

## Supporting information

Table S3 related to Figures 2-3

Table S4 related to Figure 4

## List of abbreviations

COG: conserved Oligomeric Golgi
CCD: COG Complex Dependent
vIP: Vesicle Immunoprecipitation
CCD-vIP: CCD-Vesicle Immunoprecipitation

## Authors Contributions

Farhana Taher Sumya wrote the article and made substantial contributions to conception and design, acquisition of data, analysis, and interpretation of data. Walter Saul Aragon Ramirez participated in drafting the article, performed DNA cloning and mitochondrial relocalization experiments, and interpreted the data. Vladimir V. Lupashin edited the article and made substantial contributions to conception, design and data analysis.

## Acknowledgements

We are thankful to Daniel Ungar, Rainer Duden, Elizabeth Sztul, Graham Warren as well as others who provided reagents. We are thankful to Brian Shank, Irina D. Pokrovskaya for excellent technical support and Amrita Khakurel for critical discussion. We would also like to thank the UAMS IDeA National Resource for Quantitative Proteomics (NIH/NIGMS grant R24GM137786), Digital Microscopy, sequencing and flow cytometry core facilities for the use of their facilities and expertise.

## Figure Legends

**Supplementary Figure 1. Immunoisolation of Golgi-derived vesicles. A)** Schematic diagram illustrating immune-isolation of vesicles from RPE1 COG4 KO cell line expressing COG4-mAID-mCherry and OsTIR1-9myc (RPE1 COG4-mAID) using STX5, Giantin, VAMP7 and GS15 antibodies before and after 2 hours of IAA treatment for Mass Spectrometric Analysis. The vesicles isolated from control cells are named G-vIP, V7-vIP, and GS15-vIP, and the vesicles isolated from acute COG-depleted cells are called G-CCDvIP, V7-CCDvIP, and GS15-vIP, respectively. **B)** WB analysis of Giantin, VAMP7, GS15 in Golgi fraction (P30), a soluble fraction (S30), Giantin, VAMP7, and GS15 pull-down vesicles.

**Supplementary Figure 2. Composition of different populations of vesicle proteome isolated from control cells. A)** Gene Ontology (GO) term enrichment analysis of biological process and cellular components for proteins enriched (≥2-fold, P ≤0.05) in **A)** Giantin pull-down vesicles (G-vIP), **B)** VAMP7 pull-down vesicles (V7-vIP) and **C)** GS15 pull-down vesicles (GS15-vIP) isolated from control cell compared to input (S30). **D)** Venn diagram depicting the overlap of top 100 enriched transmembrane proteins in Giantin (G-vIP), VAMP7 (V7-vIP), and GS15 pull-down vesicles (GS15-vIP) in steady state (control cell).

**Supplementary Figure 3. Composition of different populations of vesicle proteome isolated from acute COG-depleted cells. A)** Gene Ontology (GO) term enrichment analysis of biological process and cellular components for proteins enriched (≥2-fold, P ≤0.05) in **A)** Giantin pull-down vesicles (G-vIP), **B)** VAMP7 pull-down vesicles (V7-vIP) and **C)** GS15 pull-down vesicles (GS15-vIP) isolated from acute COG4 depleted cell compared to input (S30). **D)** Venn diagram depicting the overlap of top 100 enriched transmembrane proteins in Giantin (G-CCDvIP), VAMP7 (V7-CCDvIP), and GS15 pull-down vesicles (GS15-CCDvIP) in acute COG4 depleted cells.

**Supplementary Figure 4. The CCD vesicles comprise different types of Golgi resident proteins. A)** The captured image from Table 2 indicates the upregulation (Red), downregulation (Blue), and unchanged portion (transparent) of Golgi resident proteins in CCD vesicles pull-down by Giantin (G-CCDvIP), VAMP7 (V7-CCDvIP), and GS15 (GS15-CCDvIP) compared to control. The bar graph shows the percentage of upregulated, unchanged, and downregulated **B)** glycosylation machinery, **C)** other transmembrane proteins **D)** lumenal secretory proteins, and **E)** peripheral membrane proteins in G-CCDvIP, V7-CCDvIP, and GS15-CCDvIP compared to G-vIP, V7-vIP, and GS15-vIP, respectively.

**Supplementary Figure 5. Characterization of the different populations of vesicle proteome isolated from control and acute COG4-depleted cells.** Label-free mass spectrometry analysis of G-vIP/G-CCDvIP, V7-vIP/V7-CCDvIP, and GS15-vIP/GS15-CCDvIP respectively. **A)** Volcano plot depicting fold changes of proteins from four independent experiments of a direct comparison of G-vIP (control cells) and G-CCDvIP (acute COG4 depleted cells). **B)** The left panel shows the WB analysis of Giantin, MGAT1, GALNT2, and B4GALT1 in Giantin pull-down vesicles before and after IAA treatment. The right panel shows the quantification of fold enrichment of MGAT1, GALNT2, and B4GALT1 in G-CCDvIP compared to G-vIP. **C)** Volcano plot depicting results from four independent experiments of a direct comparison of V7-vIP (control cells) and V7-CCDvIP (acute COG4 depleted cells). **D)** The left panel shows the WB analysis of VAMP7, MGAT1, GALNT2, and B4GALT1 in VAMP7 pull-down vesicles before and after IAA treatment. The right panel shows the quantification of fold enrichment of MGAT1, GALNT2, and B4GALT1 in V7-CCDvIP compared to V7-vIP. **E)** Volcano plot depicting results from four independent experiments of a direct comparison of GS15-vIP (control cells) and GS15-CCDvIP (acute COG4 depleted cells). **F)** WB analysis of GS15, MGAT1, GALNT2, and B4GALT1 in GS15 pull-down vesicles before and after IAA treatment. The upper right panel shows the quantification of MGAT1, GALNT2, and B4GALT1 fold enrichment in GS15-CCDvIP compared to GS15-vIP. Volcano plots represents data from four replicates each. Colors indicate red, fold-change >2 and p < 0.05; grey, fold-change >2 or <2 and p >0.05; blue, fold-change <2 and p <0.05. Here, Red, Grey and Blue indicate significant increase, no change and significant decrease, respectively.

**Supplementary Figure 6. Comparison of CCD vesicle proteomes with RPE1 secretome. A)** Venn diagram depicting the overlap between proteins significantly enriched in G-CCDvIP, g84-CCDvIP, GS15-CCDvIP and the secretome of RPE1 cells. **B)** Venn diagram depicting the overlap between proteins significantly enriched in G-CCDvIP, V7-CCDvIP, GS15-CCDvIP and the secretome of RPE1 cells. **C)** Venn diagram depicting the overlap between soluble secretory proteins enriched in CCD-vIP of experiment 1 (EXP1) and experiment 2 (EXP2). **D)** List of soluble secretory proteins enriched in CCDvIP.

**Supplementary Figure 7. Comparison of CCD vesicle proteomes with COPI vesicle and Golgin-97 captured vesicle proteome.** A) Venn diagram depicting the overlap between Golgi proteins significantly enriched in CCDvIP and COPI proteome (HeLa cell line). B) Venn diagram depicting the overlap between Golgi proteins significantly enriched in CCDvIP and COPI essential proteome (common for three different cell lines). C) Venn diagram depicting the overlap between proteins significantly enriched in CCDvIP and Golgin-97 captured vesicles. D) Venn diagram depicting the overlap between proteins significantly enriched in GS15-CCDvIP and Golgin-97 captured vesicles.

