## Supplementary Figures 1-7 for "Comprehensive Proteomic Characterization of the Intra-Golgi Trafficking Intermediates"

A

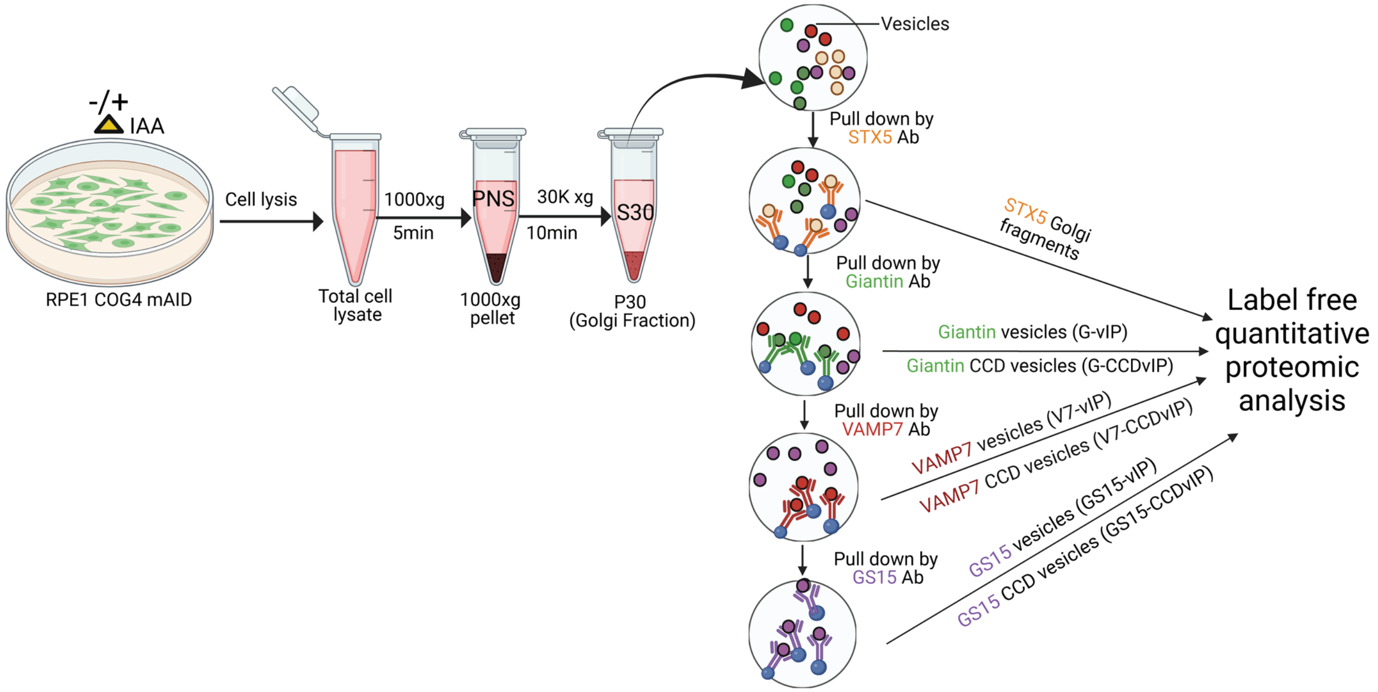

-/+
IAA
RPE1 COG4 mAID
Cell lysis
Total cell lysate
1000xg
5min
PNS
1000xg pellet
30K xg
10min
S30
P30 (Golgi Fraction)
Vesicles
Pull down by STX5 Ab
Pull down by Giantin Ab
Pull down by VAMP7 Ab
Pull down by GS15 Ab
STX5 Golgi fragments
Giantin vesicles (G-vIP)
Giantin CCD vesicles (G-CCDvIP)
VAMP7 vesicles (V7-vIP)
VAMP7 CCD vesicles (V7-CCDvIP)
GS15 vesicles (GS15-vIP)
GS15 CCD vesicles (GS15-CCDvIP)
Label free quantitative proteomic analysis


B

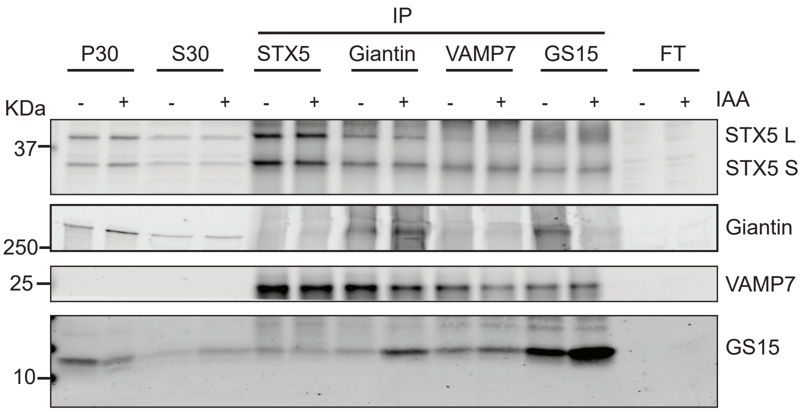

IP
P30
S30
STX5
Giantin
VAMP7
GS15
FT
KDa
- + - + - + - + - + - + - +
IAA
37
STX5 L
STX5 S
Giantin
250
25
VAMP7
GS15
10

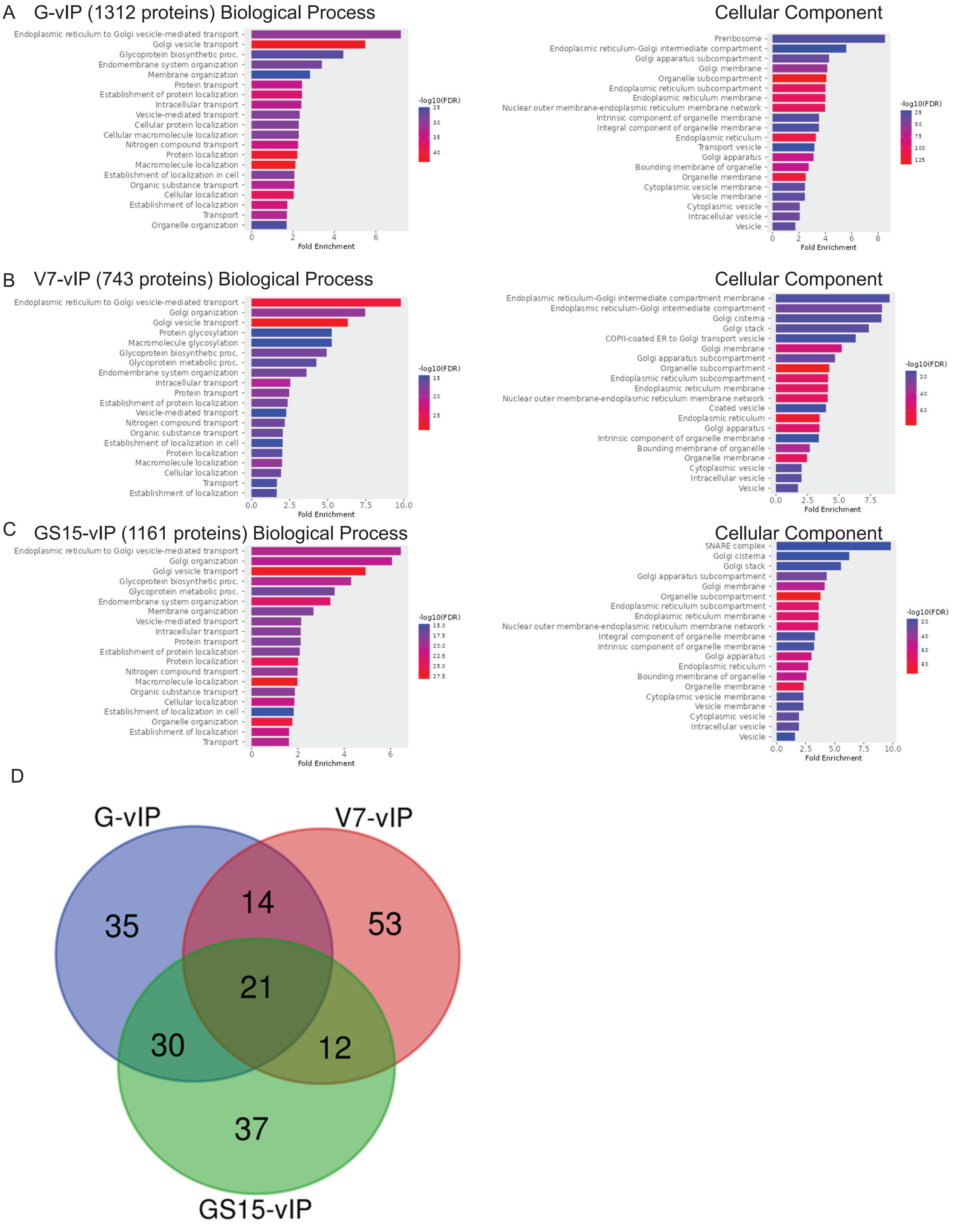
A
G-vIP (1312 proteins) Biological Process
Endoplasmic reticulum to Golgi vesicle-mediated transport
Golgi vesicle transport
Glycoprotein biosynthetic proc.
Endomembrane system organization
Membrane organization
Protein transport
Establishment of protein localization
Intracellular transport
Vesicle-mediated transport
Cellular protein localization
Cellular macromolecule localization
Nitrogen compound transport
Protein localization
Macromolecule localization
Establishment of localization in cell
Organic substance transport
Cellular localization
Establishment of localization
Transport
Organelle organization
Fold Enrichment
-log10(FDR)
Cellular Component
Preribosome
Endoplasmic reticulum-Golgi intermediate compartment
Golgi apparatus subcompartment
Golgi membrane
Organelle subcompartment
Endoplasmic reticulum subcompartment
Endoplasmic reticulum membrane
Nuclear outer membrane-endoplasmic reticulum membrane network
Intrinsic component of organelle membrane
Integral component of organelle membrane
Endoplasmic reticulum
Transport vesicle
Golgi apparatus
Bounding membrane of organelle
Organelle membrane
Cytoplasmic vesicle membrane
Vesicle membrane
Cytoplasmic vesicle
Intracellular vesicle
Vesicle
Fold Enrichment
-log10(FDR)
B
V7-vIP (743 proteins) Biological Process
Endoplasmic reticulum to Golgi vesicle-mediated transport
Golgi organization
Golgi vesicle transport
Protein glycosylation
Macromolecule glycosylation
Glycoprotein biosynthetic proc.
Glycoprotein metabolic proc.
Endomembrane system organization
Intracellular transport
Protein transport
Establishment of protein localization
Vesicle-mediated transport
Nitrogen compound transport
Organic substance transport
Establishment of localization in cell
Protein localization
Macromolecule localization
Cellular localization
Transport
Establishment of localization
Fold Enrichment
-log10(FDR)
Cellular Component
Endoplasmic reticulum-Golgi intermediate compartment membrane
Endoplasmic reticulum-Golgi intermediate compartment
Golgi cisterna
Golgi stack
COPII-coated ER to Golgi transport vesicle
Golgi membrane
Golgi apparatus subcompartment
Organelle subcompartment
Endoplasmic reticulum subcompartment
Endoplasmic reticulum membrane
Nuclear outer membrane-endoplasmic reticulum membrane network
Coated vesicle
Endoplasmic reticulum
Golgi apparatus
Intrinsic component of organelle membrane
Bounding membrane of organelle
Organelle membrane
Cytoplasmic vesicle
Intracellular vesicle
Vesicle
Fold Enrichment
-log10(FDR)
C
GS15-vIP (1161 proteins) Biological Process
Endoplasmic reticulum to Golgi vesicle-mediated transport
Golgi organization
Golgi vesicle transport
Glycoprotein biosynthetic proc.
Glycoprotein metabolic proc.
Endomembrane system organization
Membrane organization
Vesicle-mediated transport
Intracellular transport
Protein transport
Establishment of protein localization
Protein localization
Nitrogen compound transport
Macromolecule localization
Organic substance transport
Cellular localization
Establishment of localization in cell
Organelle organization
Establishment of localization
Transport
Fold Enrichment
-log10(FDR)
Cellular Component
SNARE complex
Golgi cisterna
Golgi stack
Golgi apparatus subcompartment
Golgi membrane
Organelle subcompartment
Endoplasmic reticulum subcompartment
Endoplasmic reticulum membrane
Nuclear outer membrane-endoplasmic reticulum membrane network
Integral component of organelle membrane
Intrinsic component of organelle membrane
Golgi apparatus
Endoplasmic reticulum
Bounding membrane of organelle
Organelle membrane
Cytoplasmic vesicle membrane
Vesicle membrane
Cytoplasmic vesicle
Intracellular vesicle
Vesicle
Fold Enrichment
-log10(FDR)
D
G-vIP
V7-vIP
GS15-vIP
35
14
53
21
30
12
37

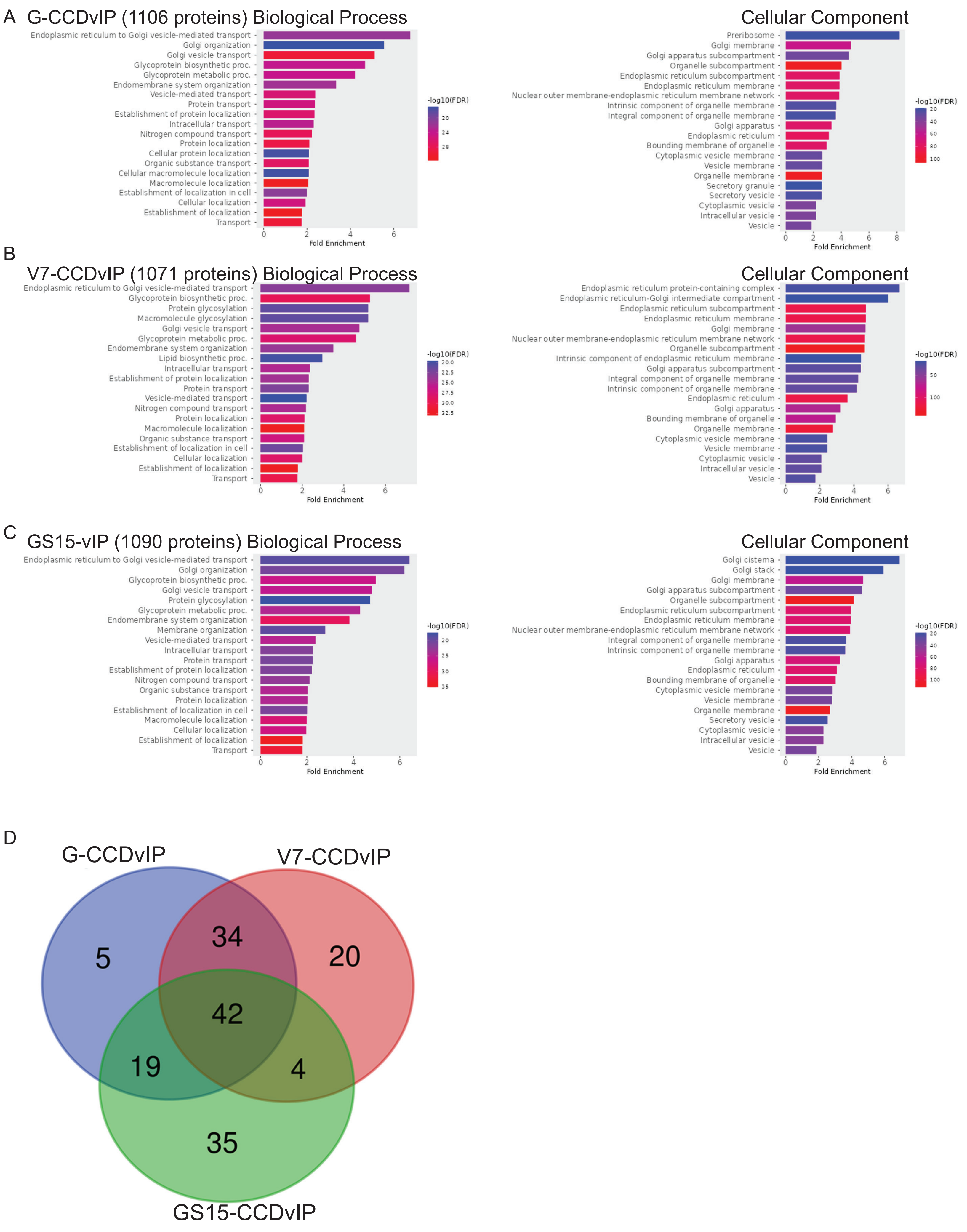
A
G-CCDvIP (1106 proteins) Biological Process
Endoplasmic reticulum to Golgi vesicle-mediated transport
Golgi organization
Golgi vesicle transport
Glycoprotein biosynthetic proc.
Glycoprotein metabolic proc.
Endomembrane system organization
Vesicle-mediated transport
Protein transport
Establishment of protein localization
Intracellular transport
Nitrogen compound transport
Protein localization
Cellular protein localization
Organic substance transport
Cellular macromolecule localization
Macromolecule localization
Establishment of localization in cell
Cellular localization
Establishment of localization
Transport
Fold Enrichment
-log10(FDR)
Cellular Component
Preribosome
Golgi membrane
Golgi apparatus subcompartment
Organelle subcompartment
Endoplasmic reticulum subcompartment
Endoplasmic reticulum membrane
Nuclear outer membrane-endoplasmic reticulum membrane network
Intrinsic component of organelle membrane
Integral component of organelle membrane
Golgi apparatus
Endoplasmic reticulum
Bounding membrane of organelle
Cytoplasmic vesicle membrane
Vesicle membrane
Organelle membrane
Secretory granule
Secretory vesicle
Cytoplasmic vesicle
Intracellular vesicle
Vesicle
Fold Enrichment
-log10(FDR)
B
V7-CCDvIP (1071 proteins) Biological Process
Endoplasmic reticulum to Golgi vesicle-mediated transport
Glycoprotein biosynthetic proc.
Protein glycosylation
Macromolecule glycosylation
Golgi vesicle transport
Glycoprotein metabolic proc.
Endomembrane system organization
Lipid biosynthetic proc.
Intracellular transport
Establishment of protein localization
Protein transport
Vesicle-mediated transport
Nitrogen compound transport
Protein localization
Macromolecule localization
Organic substance transport
Establishment of localization in cell
Cellular localization
Establishment of localization
Transport
Fold Enrichment
-log10(FDR)
Cellular Component
Endoplasmic reticulum protein-containing complex
Endoplasmic reticulum-Golgi intermediate compartment
Endoplasmic reticulum subcompartment
Endoplasmic reticulum membrane
Golgi membrane
Nuclear outer membrane-endoplasmic reticulum membrane network
Organelle subcompartment
Intrinsic component of endoplasmic reticulum membrane
Golgi apparatus subcompartment
Integral component of organelle membrane
Intrinsic component of organelle membrane
Endoplasmic reticulum
Golgi apparatus
Bounding membrane of organelle
Organelle membrane
Cytoplasmic vesicle membrane
Vesicle membrane
Cytoplasmic vesicle
Intracellular vesicle
Vesicle
Fold Enrichment
-log10(FDR)
C
GS15-vIP (1090 proteins) Biological Process
Endoplasmic reticulum to Golgi vesicle-mediated transport
Golgi organization
Glycoprotein biosynthetic proc.
Golgi vesicle transport
Protein glycosylation
Glycoprotein metabolic proc.
Endomembrane system organization
Membrane organization
Vesicle-mediated transport
Intracellular transport
Protein transport
Establishment of protein localization
Nitrogen compound transport
Organic substance transport
Protein localization
Establishment of localization in cell
Macromolecule localization
Cellular localization
Establishment of localization
Transport
Fold Enrichment
-log10(FDR)
Cellular Component
Golgi cisterna
Golgi stack
Golgi membrane
Golgi apparatus subcompartment
Organelle subcompartment
Endoplasmic reticulum subcompartment
Endoplasmic reticulum membrane
Nuclear outer membrane-endoplasmic reticulum membrane network
Integral component of organelle membrane
Intrinsic component of organelle membrane
Golgi apparatus
Endoplasmic reticulum
Bounding membrane of organelle
Cytoplasmic vesicle membrane
Vesicle membrane
Organelle membrane
Secretory vesicle
Cytoplasmic vesicle
Intracellular vesicle
Vesicle
Fold Enrichment
-log10(FDR)
D
G-CCDvIP
V7-CCDvIP
5
34
20
42
19
4
35
GS15-CCDvIP

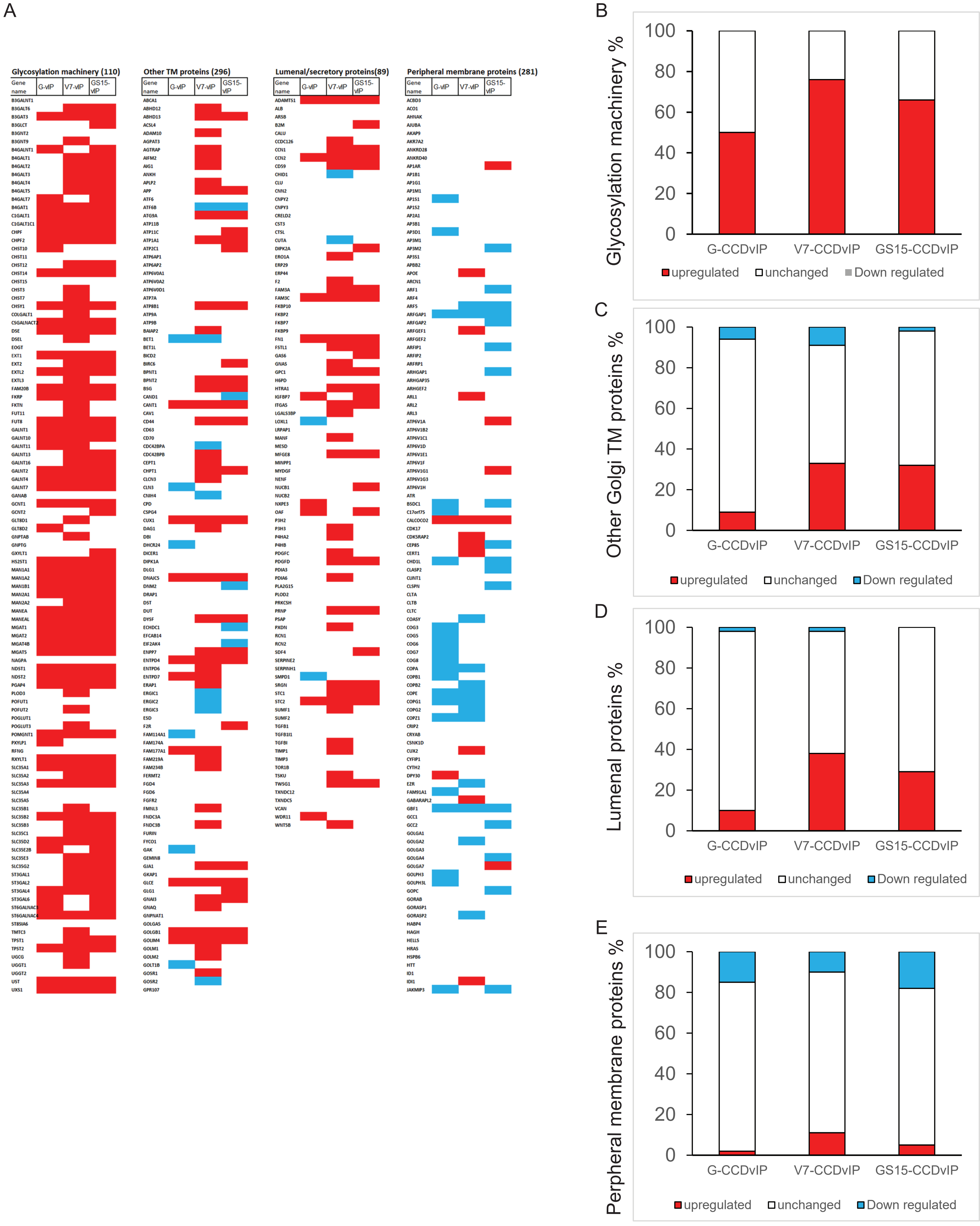
A
Glycosylation machinery (110)
Other TM proteins (296)
Lumenal/secretory proteins(89)
Peripheral membrane proteins (281)
Gene name
G-vIP
V7-vIP
GS15-vIP
B
Glycosylation machinery %
C
Other Golgi TM proteins %
D
Lumenal proteins %
E
Perpheral membrane proteins %
100
80
60
40
20
0
G-CCDvIP
V7-CCDvIP
GS15-CCDvIP
upregulated
unchanged
Down regulated

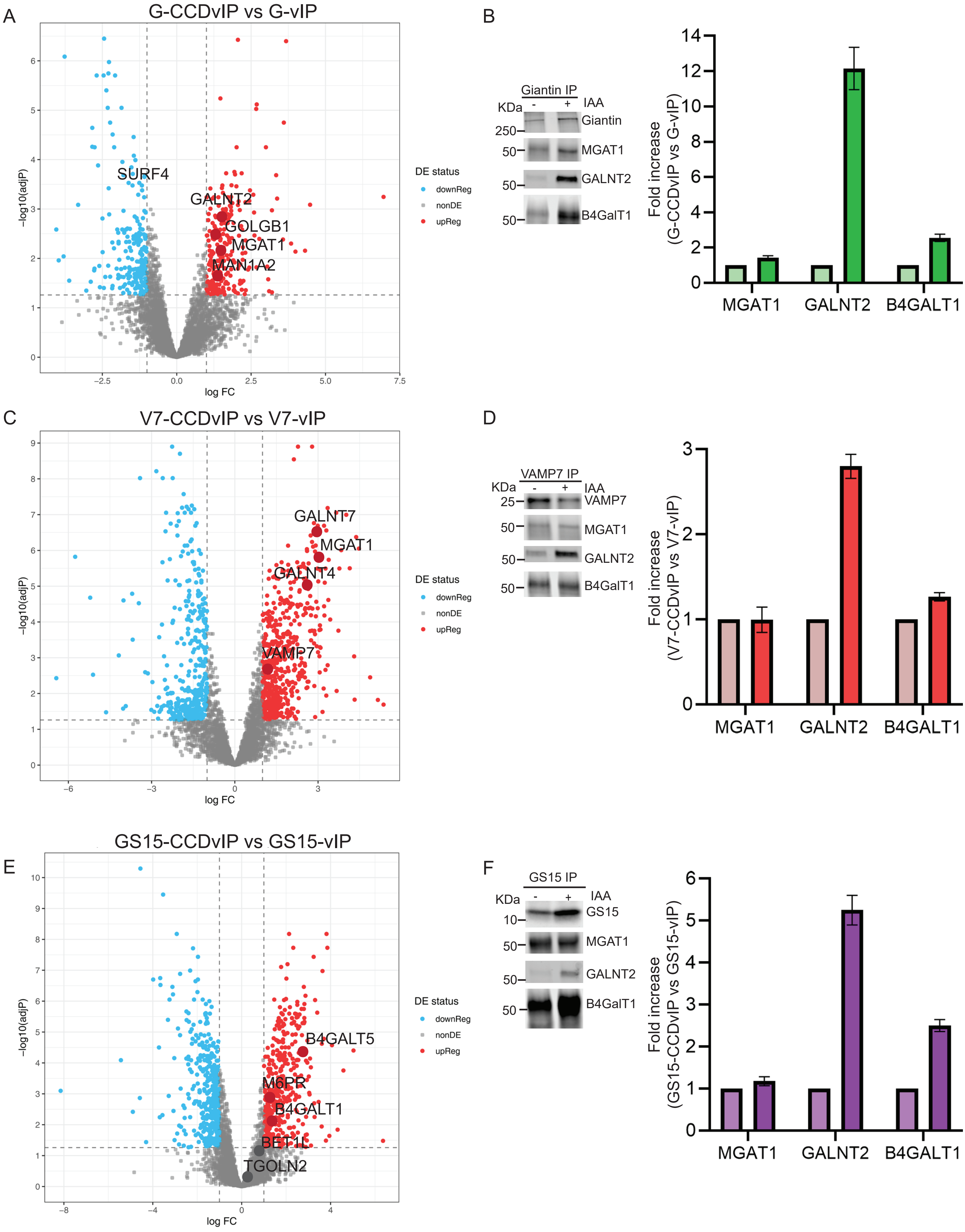
A
G-CCDvIP vs G-vIP
SURF4
GALNT2
GOLGB1
MGAT1
MAN1A2
−log10(adjP)
log FC
DE status
downReg
nonDE
upReg
B
Giantin IP
KDa
IAA
Giantin
MGAT1
GALNT2
B4GalT1
250
50
Fold increase (G-CCDvIP vs G-vIP)
MGAT1
GALNT2
B4GALT1
C
V7-CCDvIP vs V7-vIP
GALNT7
MGAT1
GALNT4
VAMP7
−log10(adjP)
log FC
DE status
downReg
nonDE
upReg
D
VAMP7 IP
KDa
IAA
VAMP7
MGAT1
GALNT2
B4GalT1
25
50
Fold increase (V7-CCDvIP vs V7-vIP)
MGAT1
GALNT2
B4GALT1
E
GS15-CCDvIP vs GS15-vIP
B4GALT5
M6PR
B4GALT1
BET1L
TGOLN2
−log10(adjP)
log FC
DE status
downReg
nonDE
upReg
F
GS15 IP
KDa
IAA
GS15
MGAT1
GALNT2
B4GalT1
10
50
Fold increase (GS15-CCDvIP vs GS15-vIP)
MGAT1
GALNT2
B4GALT1

A

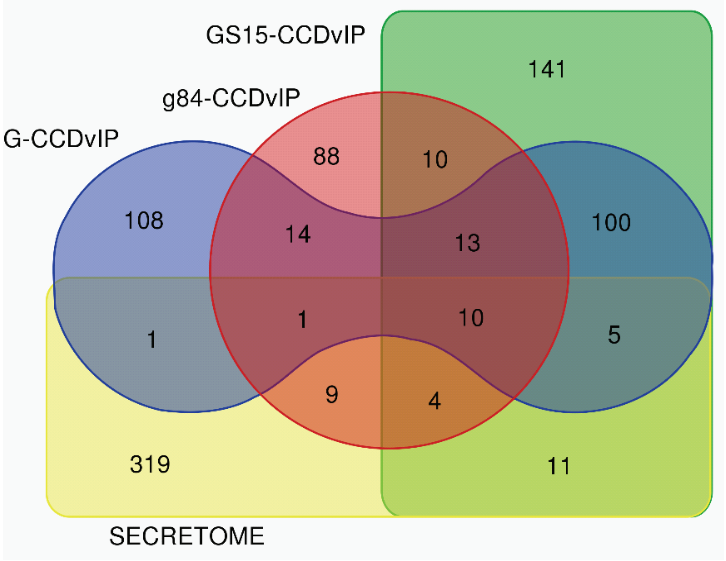


B

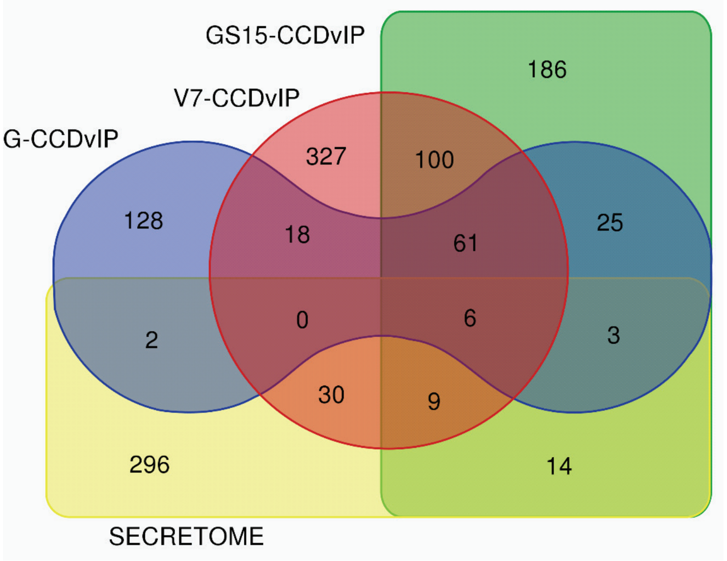


C

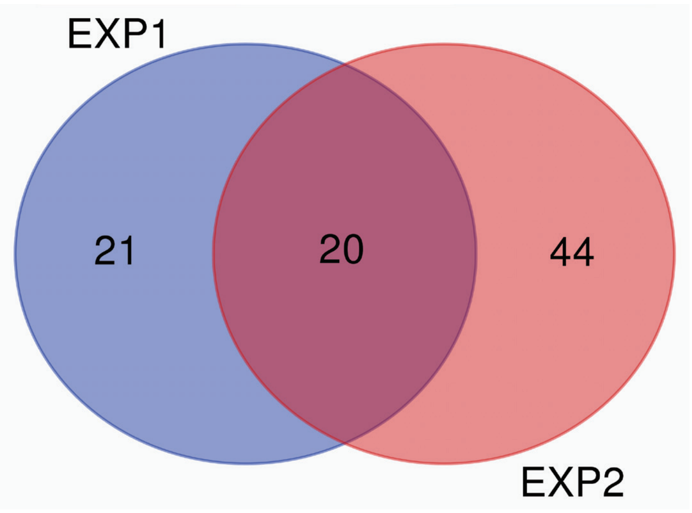


D

| Names | total | elements |
|---|---|---|
| EXP1 SECRETOME EXP2 SECRETOME | 20 | CFB SERPINE1 LAMB1 FAM3C PTX3 PXDN P4HA2 SDF4 LOX NT5E NUCB1 THBS1 STC2 CD59 TGFBI PLOD3 TIMP1 LGALS3BP IGFBP7 DCD |
| EXP1 SECRETOME | 21 | SOD3 HSP90B1 SERPINE2 C4A CPA4 NUCB2 DKK3 APOA1 TXNDC5 COL1A1 FKBP2 RCN1 SMPD1 GANAB ECM1 PDIA4 SERPINA3 TXNDC12 CALR PLOD2 CTSC |
| EXP2 SECRETOME | 44 | FSTL1 THY1 PRSS23 PRDX4 SPARC INHBA P4HA1 MFGE8 HYOU1 SFRP1 MANF FUCA1 PLBD2 TGFB2 PRNP WNT5B LAMC1 HTRA1 GNS MATN2 B2M NXPE3 TWSG1 CRTAP GPC1 PDGFC HSPA13 COL12A1 GSN OAF DNAJC3 PDIA6 PDGFD GAS6 FST FN1 TOR1A NPC2 DNAJB11 COLGALT1 SEMA7A SPOCK1 PLOD1 PLAUR |

RED – SECRETORY
BLUE – GOLGI
GREEN – ER
ORANGE – GPI ANCHORED

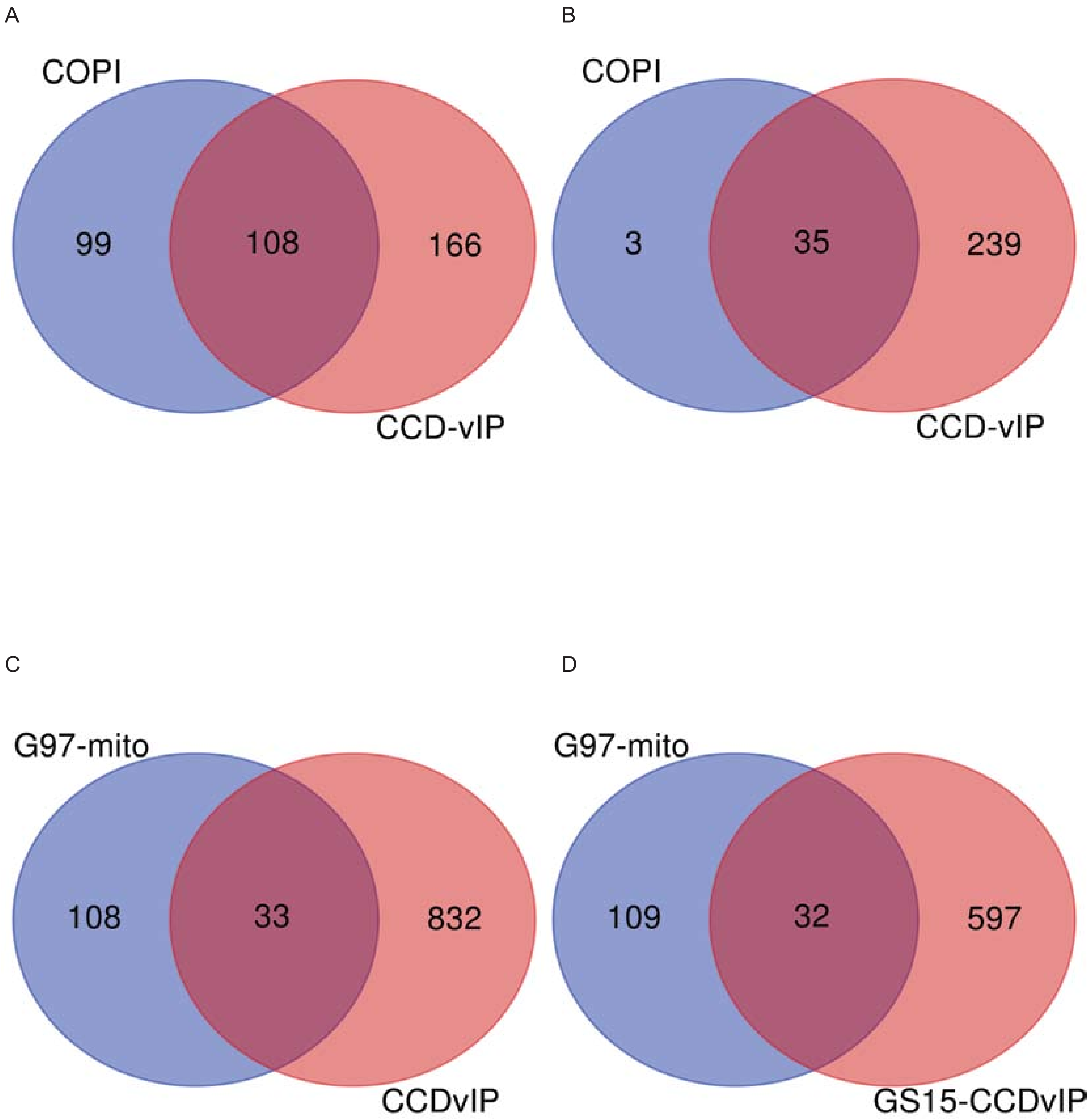
A
COPI
99
108
166
CCD-vIP
B
COPI
3
35
239
CCD-vIP
C
G97-mito
108
33
832
CCDvIP
D
G97-mito
109
32
597
GS15-CCDvIP
